# A new method for enamel amino acid racemization dating: a closed system approach

**DOI:** 10.1101/453340

**Authors:** Marc R. Dickinson, Adrian M. Lister, Kirsty E. H. Penkman

**Affiliations:** Department of Chemistry, University of York, York, YO10 5DD, UK; Natural History Museum, Cromwell Road, London, SW7 5BD, UK

**Keywords:** Enamel, amino acid racemization, closed system, biphasic separation, geochronology, IcPD Highlights

## Abstract

Analysis of the predictable breakdown of proteins and amino acids in ancient biominerals enables age estimation over the Quaternary. We postulate that enamel is a suitable biomineral for the long-term survival of endogenous amino acids. Analysis of multiple amino acids for geochronological studies is typically achieved using a RP-HPLC method. However, the low concentrations of amino acids coupled with high concentrations of inorganic species make accurate determination of amino concentrations challenging. We have developed a method for the routine preparation of multiple enamel samples using biphasic separation. Furthermore, we have shown that amino acids that exhibit effectively closed system behaviour can be isolated from enamel through an exposure time of 72 h to bleach. Elevated temperature experiments investigating the processes of intra-crystalline protein degradation (IcPD) do not appear to match the patterns from fossil samples, reinforcing the need for a comprehensive understanding of the underlying mechanisms of protein degradation. This novel preparative method isolates intra-crystalline amino acids suitable for the development of mammalian geochronologies based on enamel protein degradation. The lower rates of racemisation in enamel (cf. *Bithynia* opercula) suggest that the enamel AAR may be able to be used as a relative dating technique over time scales > 2.8 Ma. Enamel AAR has the potential to estimate the age of mammalian remains past the limit of all other current direct dating methods, providing an invaluable tool for geochronological studies.

---

- Development of a biphasic separation method for the extraction of amino acids from enamel
- Isolation of intra-crystalline amino acids in enamel
- Testing of the closed system behaviour of the enamel intra-crystalline amino acids
- Modelling the kinetic behaviour of enamel amino acids
- Comparison of enamel and *Bithynia* opercula fossil data and elevated temperature experiments
- we demonstrate promising results for the analysis of AAR in enamel for geochronological purposes

## 1 Introduction

Mammalian teeth are often found in palaeontological deposits, providing an excellent target for direct dating. However, mammalian remains older than the limits of radiocarbon analysis (~50 ka, MacPhee *et al.*, 2002; Jacobi, 2006) are challenging to date and therefore frequently rely on chronological analysis of associated material (e.g. Frouin *et al.*, 2017). Currently, electron spin resonance (ESR) and U-series (which can be used together or separately) are the only commonplace direct dating techniques available for skeletal remains older than ~50 ka and spanning the Middle to Late Pleistocene (Dirks *et al.*, 2017; Hershkovitz, *et al.*, 2018). Both these techniques require an accurate reconstruction of the U-uptake history, but as teeth (and bones) are open systems for uranium, modelling this uptake is challenging. U-uptake modelling (such as diffusive-absorption models) can, however, provide an age estimation up to ~500 ka on small samples on the order of ~ 10-50 mg (Dirks *et al.*, 2017; Hershkovitz, *et al.*, 2018), which can be pushed further back (~750 ka) using combined Pa/U and Th/U data. Critically, at present this is restricted by the requirement for significantly larger sample sizes of up to ~1 g (Grün *et al.*, 2010; Duval, 2015). Moreover, it is thought that the best source of material is often from the centre of the sample, thereby requiring relatively destructive sampling methods (Grün *et al.*, 2010).

Amino acid racemization (AAR) analysis has proven a valuable technique for age estimation over Quaternary timescales (~2.5 Ma) for a variety of calcium carbonate based biominerals (Wehmiller *et al.*, 2012; Hendy *et al.*, 2012; Refsnider *et al.*, 2013) and it requires comparatively small sample mass (~10 mg). AAR dating of mammalian collagen-based biominerals (e.g. bone and dentine) has had more limited success (Taylor 1983; Bischoff & Rosenbauer, 1981; Bada 1985; Blackwell *et al.*, 1990; Marshall, 1990). Amino acid kinetic studies, conducted on dentine have shown a good adherence between the temperature-induced models and fossil data for some amino acids, but importantly not all (Canoira *et al.*, 2003) and these issues are compounded by the non-linear degradation of collagen (Collins *et al.*, 2009). However, critically the concerns associated with leaching and contamination of such open systems preclude accurate dates (Towe, 1980; Hare, 1988; Bravenec *et al.*, 2018).

Early studies postulated that amino acids can become trapped during the formation of certain biominerals (Towe and Thompson, 1972); recent research has shown mineral facets ranging in size from ~2.5-38.4 nm can be found in molluscan nacre, with elemental analysis indicating a higher carbon content in these voids (Gries *et al.*, 2009). This suggests that organic material is likely to be trapped within these voids, and therefore could be isolated from the external chemical environment. Prolonged exposure of powdered mollusc shells to bleach has been shown to isolate a fraction of organic matter that is resistant to oxidative treatment (Sykes 1995, Penkman *et al.*, 2007, 2008), defined as the intra-crystalline fraction, with the potential to act as a closed system for protein degradation. As this contradicts the strictest definition of crystals (which cannot contain large macromolecules), here we define “intra-crystalline” as the fraction of organic matter that is resistant to prolonged oxidative treatment (Sykes *et al.*, 1995). Bleach has been used to isolate intra-crystalline proteins and amino acids in mollusc shells (Sykes, 1995; Penkman *et al.*, 2008; Demarchi *et al.*, 2013a), opercula (Penkman *et al.*, 2011) and coral (Hendy *et al.*, 2012, Tomiak *et al.*, 2013).

These voids imaged in nacre may also be present in tooth enamel, and expansion of AAR analysis to the routine dating of mammalian remains would be highly valuable, for example in helping to elucidate the migration and evolution of fauna in response to Pleistocene climate change. Enamel is composed of a form of hydroxyapatite (HA; calcium phosphate) which is heavily mineralised. The determination of L- and D- amino acids for geochronological purposes is typically achieved by analysis by HPLC with fluorescence detection modified from Kaufman & Manley (1998). However, the application of AAR analysis on enamel is not without challenges, as high concentrations of inorganic salts originating from demineralisation of the enamel crystal structure (calcium phosphate) result in peak response suppression, and unstable and raised baselines limiting accurate quantification (Griffin, 2006).

We therefore present a new method for the preparation of enamel for AAR analysis. Through RP-HPLC, FTIR and TEM analyses, we evaluate whether intra-crystalline amino acids and proteins are present in enamel and if they exhibit closed system behaviour. Long term degradation of enamel is simulated through elevated temperatures and compared to results from fossil material to gain a better understanding of the reaction kinetics of racemization.

To this end, a series of experiments was performed on enamel:

1. Optimisation of the biphasic separation of inorganic species from amino acids (Section 3), using:

a. FT-IR analysis of both the supernatant (containing the amino acids) and the underlying gel
b. optimising the volume of KOH used in the separation
c. testing the impact of the optimised method on the relative concentration of amino acids and their D/L values
d. evaluation of the expected analytical error
2. Optimisation of the oxidation method for the isolation of any intra-crystalline amino acids from the enamel (Section 2).
3. Elevated-temperature experiments designed to investigate potential leaching (diffusive loss) of amino acids from both the intra- and inter-crystalline fractions, as well as to study the kinetics of intra-crystalline protein degradation (IcPD; Section 5).
4. Analysis of racemization in fossil proboscidean enamel from the UK, providing a pilot relative geochronology (Section 5.5.5 - 5.5.6).

## 2 Materials and optimised methods

As some of the methods were optimised over the course of the study (Sections 3 - 4), the final optimised methods are detailed in this section.

### 2.1 Enamel samples

Twelve teeth were sampled from a range of contexts varying in age from modern to ~2.2 Ma (Table 1 and SI); fossil material was selected from sites which had independent evidence of age.

**Table 1.**
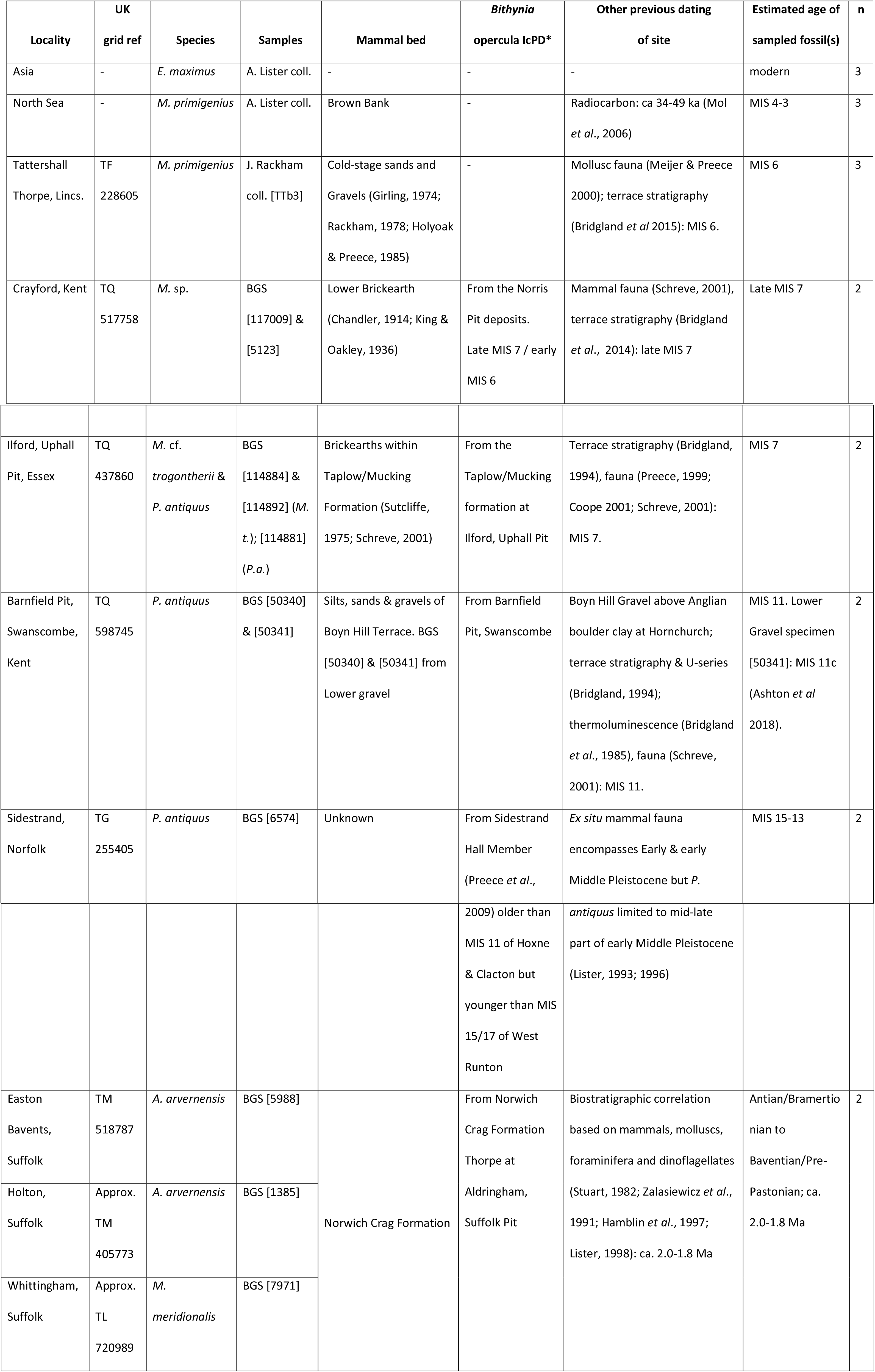
Sample information for the fossil enamel and comparative *Bithynia* opercula, taken from Penkman *et al.*, 2013. *E. = Elephas, P*. =*Palaeoloxodon, M. = Mammuthus, A. = Anancus*

#### 2.1.1 Hydroxyapatite with amino acid reference solutions

To mimic an enamel fossil sample with a known relative concentration of amino acids and D/L values, 1:1 and 1:4 D:L amino acid reference solutions (10 μL) were added to reagent grade hydroxyapatite (~2 mg; <200 nm particle size; Sigma Aldrich) and dried by centrifugal evaporation. The 1:1 amino acid reference solution mimics an older sample in which the amino acids have reached equilibrium, while the 1:4 amino acid reference solution mimics a younger sample. The reference solutions contain the free amino acids: Arg, Asp, Glu, Gly, His, Ile (D-isomer D-alle), Leu, Lys, Met, Phe, Ser, Thr, Tyr and Val with a D to L-isomer ratio of ~1 (1:1) or 0.2 (1:4) and the internal standard (L-*homo*-arginine).

**Table 2.** Samples used in each series of experiments. Optimisation and testing of the biphasic separation of inorganic species from amino acids method (OBS); Isolation of the intra-crystalline fraction (IIcF); Elevated temperature experiments to test for closed system behaviour (ET); fossil comparison to *Bithynia* opercula (F)

| Sample code | BGS number | Species | Site | Experiment |
| --- | --- | --- | --- | --- |
| MoE1 | - | *E. maximus* | - | ET |
| NSM1 | - | *M. primigenius* | North Sea | ET |
| TaM1 | - | *M. primigenius* | Tattershall Thorpe | IlcF & OBS |
| CrM1 | 117009 | *M. sp.* | Crayford | F |
| CrM2 | 5123 | *M. sp.* | Crayford | F |
| IIP1 | 114881 | *P. antiquus* | Ilford | F |
| IIM1 | 114884 | *M. trogontherii* | Ilford | F |
| IIM2 | 114892 | *M. trogontherii* | Ilford | F |
| SwP1 | 50340 | *P. antiquus* | Swanscombe | F |
| SwP2 | 50341 | *P. antiquus* | Swanscombe | F |
| SiP1 | 6574 | *P. antiquus* | Sidestrand | F |
| NCA1 | 5988 | *A. arvernensis* | Easton Bavents | F |
| NCA2 | 7971 | *A. arvernensis* | Whittingham | F |
| NCM1 | 1385 | *M. meridionalis* | Holton | F |

### 2.2 Enamel sampling

An enamel chip was removed from each tooth with a Dremel drill and the exposed outside layer removed using an abrasive rotary burr drill bit. Care was taken to remove any visible dark discolouration that was probably due to iron or manganese staining (Turner-Walker, 2007; Garot *et al.*, 2017). To eliminate powder resulting from the initial drilling, chips were sonicated for 3 min in HPLC-grade water (Sigma-Aldrich) and then ethanol (VWR, analytical-grade), and air-dried before being finely powdered using an agate pestle and mortar.

### 2.3 Optimised NaOCl procedure

Approximately 30 mg of powdered enamel was weighed into a 2 mL microcentrifuge tube (Eppendorf), and NaOCl (12%, Fisher Scientific, analytical grade, 50 µL mg^−1^ of enamel) was added. Samples were exposed to NaOCl for 72 h and were continuously agitated to ensure complete exposure. The NaOCl was removed and the powdered enamel was washed five times with HPLC-grade water. A final wash with methanol (Sigma-Aldrich, HPLC-grade) was used to remove remaining NaOCl, before the sample was left to air dry overnight.

### 2.4 Preparation of FAA and THAA fractions

Powdered enamel samples were accurately weighed into two fractions. One sample was treated for their free amino acid (FAA) content and the other their total hydrolysable amino acid (THAA) content.

#### 2.4.1 Hydrolysis

THAA samples were dissolved in HCl (7 M, 20 μL mg^−1^) and heated in a sterile sealed glass vial at 110 °C for 24 h, whilst the vials were purged with N_2_ to prevent oxidation. The acid was removed by centrifugal evaporation.

#### 2.4.2 Biphasic phosphate ion removal

THAA samples were re-dissolved in HCl (1 M, 20 μL mg^−1^) and FAA samples were demineralised in HCl (1 M, 25 μL mg^−1^) in a sterile 0.5 mL microcentrifuge tube (Eppendorf) and sonicated for 10 min or until all visible signs of undissolved material had disappeared. To remove the high concentrations of phosphate ions (thought to impact on the accuracy of the RP-HPLC analysis), an additional protocol to those outlined for the standard analysis of amino acids in calcium carbonate-based biominerals (Kaufman & Manley, 1998; Penkman *et al.*, 2008) was developed. The optimised method is as follows: KOH (1 M, 28 μL mg^−1^, Fisher Scientific, analytical grade) was added to the acidified solutions, whereupon a mono-phasic cloudy solution formed. The samples were then briefly agitated to homogenise the solution. The sample was centrifuged at 13000 rpm for 10 min causing a clear supernatant to form above a gel; the biphasic separation (Figure 1). The supernatant was extracted and dried by centrifugal evaporation.

**Figure 1.**
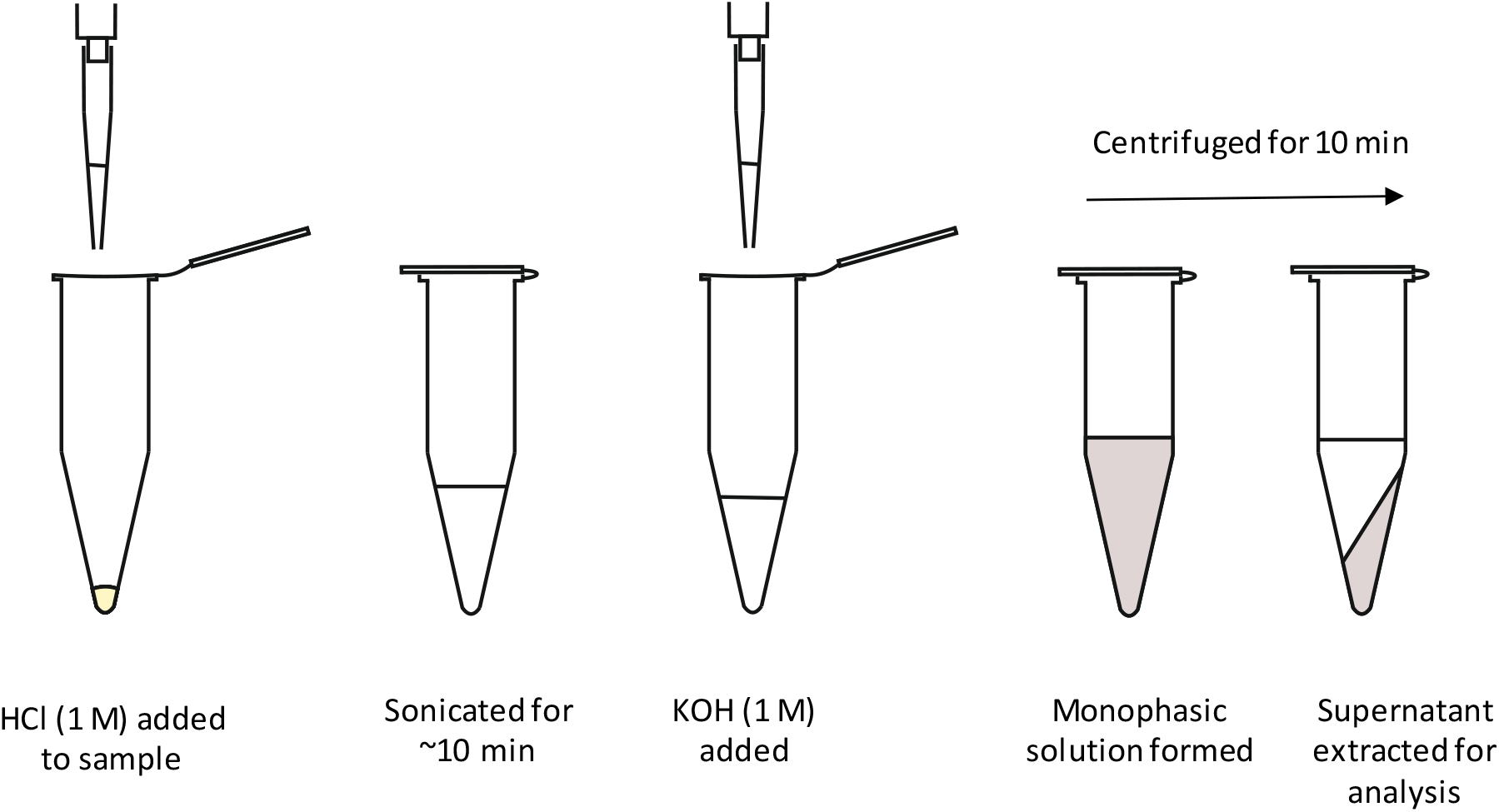
*Schematic of the biphasic extraction of amino acids, designed to reduce phosphate ion concentration in the samples undergoing RP-HPLC analysis*.

### 2.5 RP-HPLC Analysis

Samples were rehydrated with a solution containing an internal standard (L-*homo*-arginine; 0.01 mM), sodium azide (1.5 mM) and HCl (0.01 M), to enable quantification of the amino acids. Analysis of chiral amino acid pairs was achieved using an Agilent 1100 Series HPLC fitted with a HyperSil C18 base deactivated silica column (5 μm, 250 × 3 mm) and fluorescence detector, using a modified method outlined by Kaufman and Manley (1998). The column temperature was controlled at 25 °C and a tertiary system containing sodium buffer (23 mM sodium acetate trihydrate, sodium azide, 1.3 μM EDTA, adjusted to pH 6.00 ±0.01 with 10 % acetic acid and sodium hydroxide), acetonitrile and methanol was used as a solvent.

### 2.6 FT-IR analysis

FT-IR analysis of both the dried gel and supernatant were conducted so the fate of the inorganic phosphate species could be determined. Fossil mammoth enamel from Tattershall Thorpe (Table 1) was powdered and bleached according to the optimised protocol (Section 2.2–2.3). Bleached enamel was treated as if analysing FAAs using the optimised biphasic separation (Section 2.4). The gel that formed during the biphasic separation was dried by centrifugal evaporation. Both the dried gel and supernatant produced solids suitable for IR analysis: the gel formed a white powdery pellet, whist the supernatant formed transparent crystals. To identify the location of the phosphate species, FT-IR analysis was carried out on each fraction as well as on pure hydroxyapatite for reference, using a Perkin Elmer Spectrum 2 IR spectrometer set to scan from 4000 – 350 cm^−1^ with a resolution of 2 cm^−1^.

## 3 Optimisation and testing of the biphasic separation of inorganic species from amino acids

To remove the amino acids from the phosphate species to enable RP-HPLC analysis, the biphasic separation approach described above was optimised.

### 3.1 Biphasic separation optimisation methods

To develop an optimised method for the biphasic separation, three aspects were tested in detail:

a. the impact of the volume of KOH solution used in the preparation
b. the impact of the optimised method on the relative concentration of amino acids and their D/L values
c. evaluation of the expected analytical error

For part a. (testing the impact of KOH volume); fossil enamel from the Thorpe sand and gravel at Tattershall was powdered and the intra-crystalline amino acids were isolated. THAAs were prepared (Section 2.4.1) and re-dissolved in the minimum volume of HCl (1 M, 20 μL mg^−1^) and FAA samples were demineralised in HCl (1 M, 25 μL mg^−1^). Different volumes of KOH (1 M, Fisher-Scientific) were added to these sub-samples prepared for their THAA (12 - 65 μL) and FAA (12 - 180 μL) contents to test for the optimum volumes required. A monophasic cloudy solution formed upon addition of KOH. The solution was centrifuged at 13000 rpm for 10 min and a clear supernatant formed above a gel (Figure 1). The supernatant was extracted and dried by centrifugal evaporation. Most samples were run in triplicate and were analysed by RP-HPLC (Section 2.5). However, upon addition of the rehydration fluid, some samples were insoluble due to the presence of high concentrations of calcium phosphate; in these cases, only one replicate was analysed by RP-HPLC to prevent damage to the HPLC system.

For part b. (quantifying the impact identifying impact of the optimised method on amino acid composition); 1:1 and 1:4 amino acid reference solutions were added to powdered hydroxyapatite. Samples followed the optimised protocol outlined for FAA samples post bleaching (Sections 2.4.2 and 2.5).

For part c. (estimation of analytical errors); *M. primigenius* tooth enamel from Tattershall Thorpe (Table 1) was powdered and homogenised (Section 2.2). Eight sub-samples were taken and run independently for their THAA and their FAA content as described in sections 2.2 - 2.5. Each sub-sample was run in duplicate. Some of the amino acids do not have reported D/L values due to low concentrations (Ile) and/or the presence of co-eluting peaks (THAA-Tyr, Leu and Ile), making accurate quantification impossible; in both instances, it is the usually the D-isomer peak that is affected.

### 3.2 Optimising the volume of KOH added during the biphasic separation of phosphate species and amino acids: results and discussion

A biphasic method for separation of phosphate species from amino acids has been developed to eliminate the chromatographic issues caused by high concentrations of inorganic species during RP-HPLC analysis.

#### 3.2.1 FT-IR analysis of the gel and supernatant

The IR spectrum for the dried gel displays absorption bands assigned to phosphate (PO_4_^3-^): vibrational frequencies at 1000 cm^−1^ (v_3_), 962 cm^−1^ (v_1_) and 561 cm^−1^ (v_4_; Paz *et al.*, 2012), all of which are also present in the spectrum of pure hydroxyapatite (Figure 2). The IR spectrum for the supernatant does not contain any discernible absorption bands (Figure 2), indicating that it does not contain detectable concentrations of phosphate species. The crystals formed by the dried supernatant are most likely KCl, which is not detectable by IR analysis. The presence of phosphate absorption bands in the gel and not in the supernatant indicates that phosphate species are removed from solution using this biphasic extraction method.

**Figure 2.**
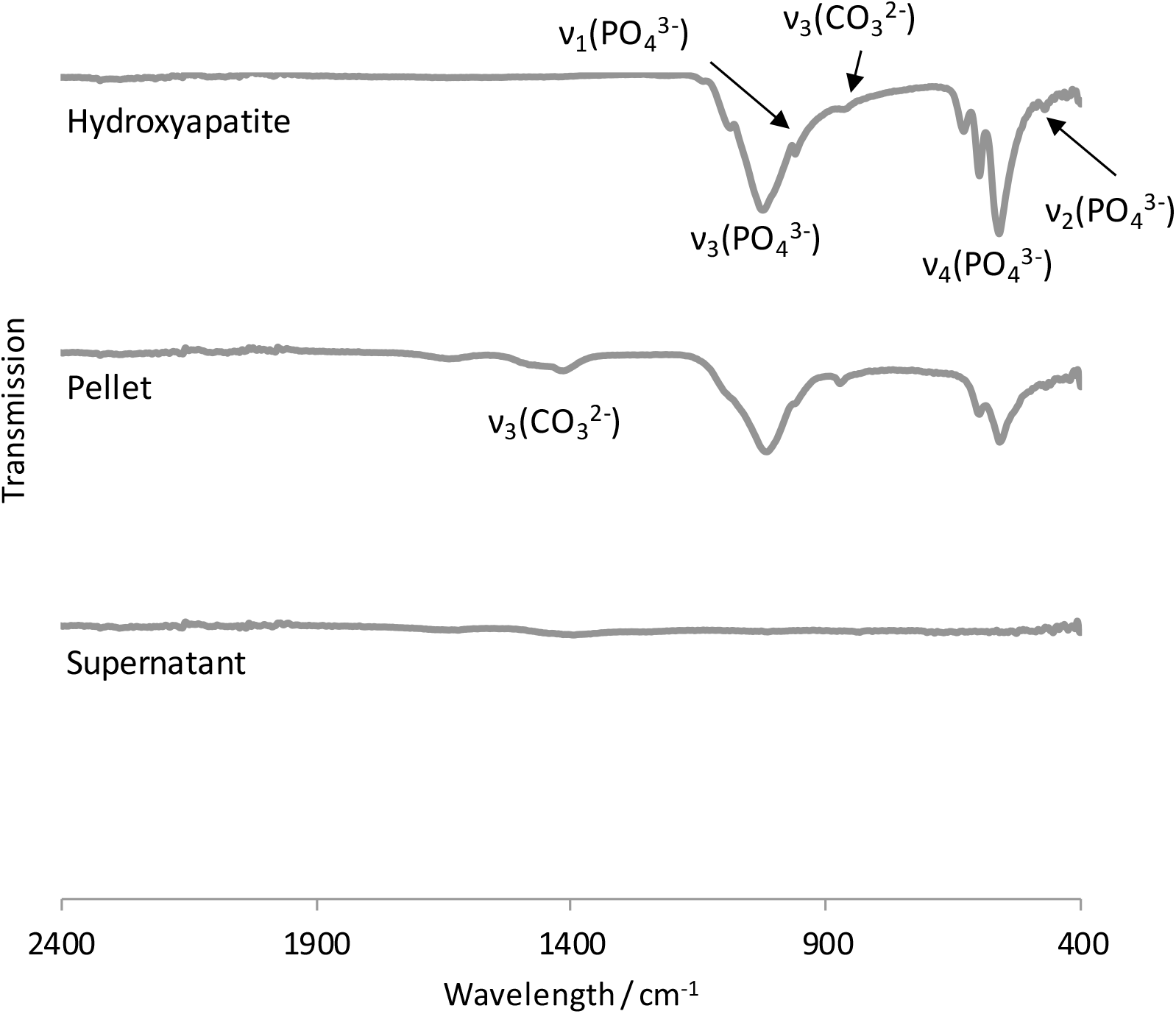
*Stacked FT-IR spectra of pure hydroxyapatite (top), the gel formed after biphasic separation (middle), exhibiting phosphate absorption bands and the supernatant extracted (bottom). Absorption bands associated with inorganic salts have been labelled based on assignment from Paz et al. (2012). This data indicates that phosphate is incorporated into the gel, with none detectable in the supernatant then used for amino acid analysis. Carbonate peaks are also present due to CO2 substitution in the hydroxyapatite crystal structure (Paz *et al.*, 2012)*.

#### 3.2.2 Changes in FAA and THAA concentration with increasing KOH volume

For the samples analysed for their FAAs, the measured concentration of Asx, Glx and Ala initially drops with increasing KOH volume (12-20 μL; Figure 3); this is most likely due to an increasing HPLC response to the internal standard, as the inorganic phosphates are more effectively removed and peak suppression was reduced. Above 16 µL mg^−1^ of KOH, the internal standard peak response lies within the expected range (i.e. of the rehydration solution alone), but peak area reduces again as excess KOH (> 40 µL mg^−1^) start to cause peak suppression. The measured concentration of Asx when 180 μL of KOH solution was added is higher than at lower volumes of KOH, but this is not replicated in the other amino acids (Figure 3). The LhArg peak response at this concentration is lower than at other concentrations, indicating that Asx has a different response to the higher levels of KOH than the other amino acids (including LhArg). This might lead to inaccuracies in analysing the relative amino acid composition of fossil samples. Additionally, the samples analysed at the highest and lowest volumes of KOH solution (12, 120 and 180 µL mg^−1^) induced cloudy solutions prior to HPLC injection, suggesting higher concentrations of inorganic salts. Volumes of 20-65 µL mg^−1^ of KOH used to induce gel formation are therefore most likely to yield reliable amino acid data, and a value of 28 µL mg^−1^ is recommended.

**Figure 3.**
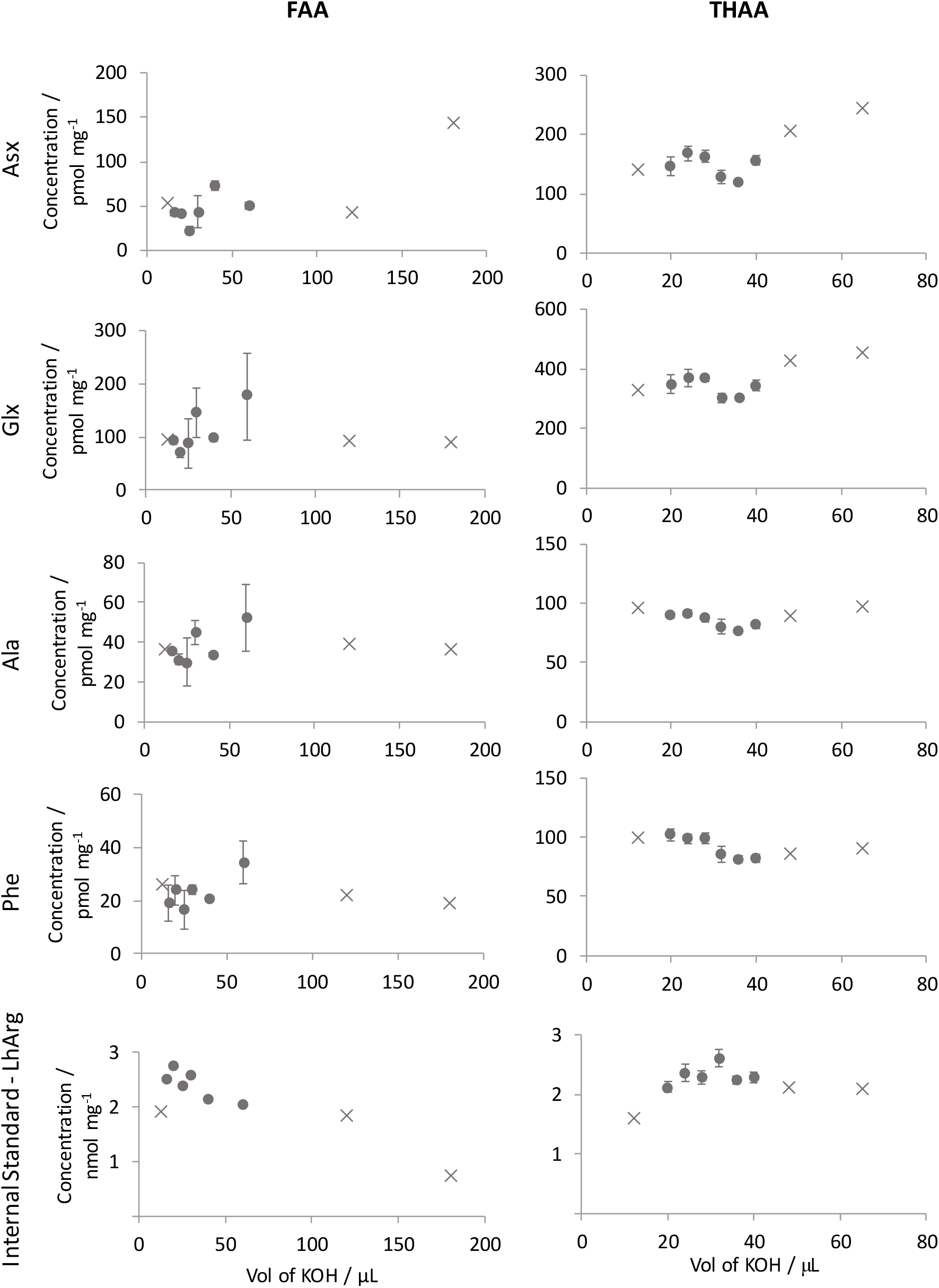
*FAA (Left) and THAA (right) biphasic separation optimisation: Change in concentration of amino acids when changing the volume of KOH added (top four rows), the change in the HPLC response to the internal standard when changing the volume of KOH (bottom panel). Error bars depict one standard deviation about the mean (n = 3). Samples that were cloudy prior to HPLC injection have been plotted as crosses, only one sample of each of these volumes were analysed*.

Generally, a higher precision (cf. 1σ error bars Figure 3) was observed for the THAA measured concentrations than for comparable FAA fractions (Figure 3) and this is most likely due to the higher amounts of amino acids in the THAA fractions, making quantification of integrated HPLC peak areas more precise. The measured concentration for Glx, Ala, and Phe are relatively stable with increasing KOH volume (Figure 3). In contrast, the measured concentration of Asx increases at KOH volumes greater than 42 μL, whilst the LhArg peak area response remains relatively stable. The change in Asx concentration with KOH volume alludes to either a bias in the peak suppression when compared to other amino acids (including the internal standard), or that the concentration of Asx could be biased by gel formation (see Section 3.3).

#### 3.2.3 Changes in D/L value with increasing KOH volume

The D/L value for most amino acids is unaffected by changes in KOH volume, except for Ala prepared with 12 μL KOH and the THAA Glx and Phe prepared with 70 μL KOH (Figure 4). This suggests the biphasic gel formation does not bias the D/L values at the intermediate volumes (FAA: 16 - 120 μL; THAA: 20 - 65 μL). Whilst a range of volumes of KOH may be appropriate, a standard volume of 28 μL mg^−1^ has been selected for the biphasic separation of amino acids from phosphate species, as this volume is unaffected by peak suppression.

**Figure 4.**
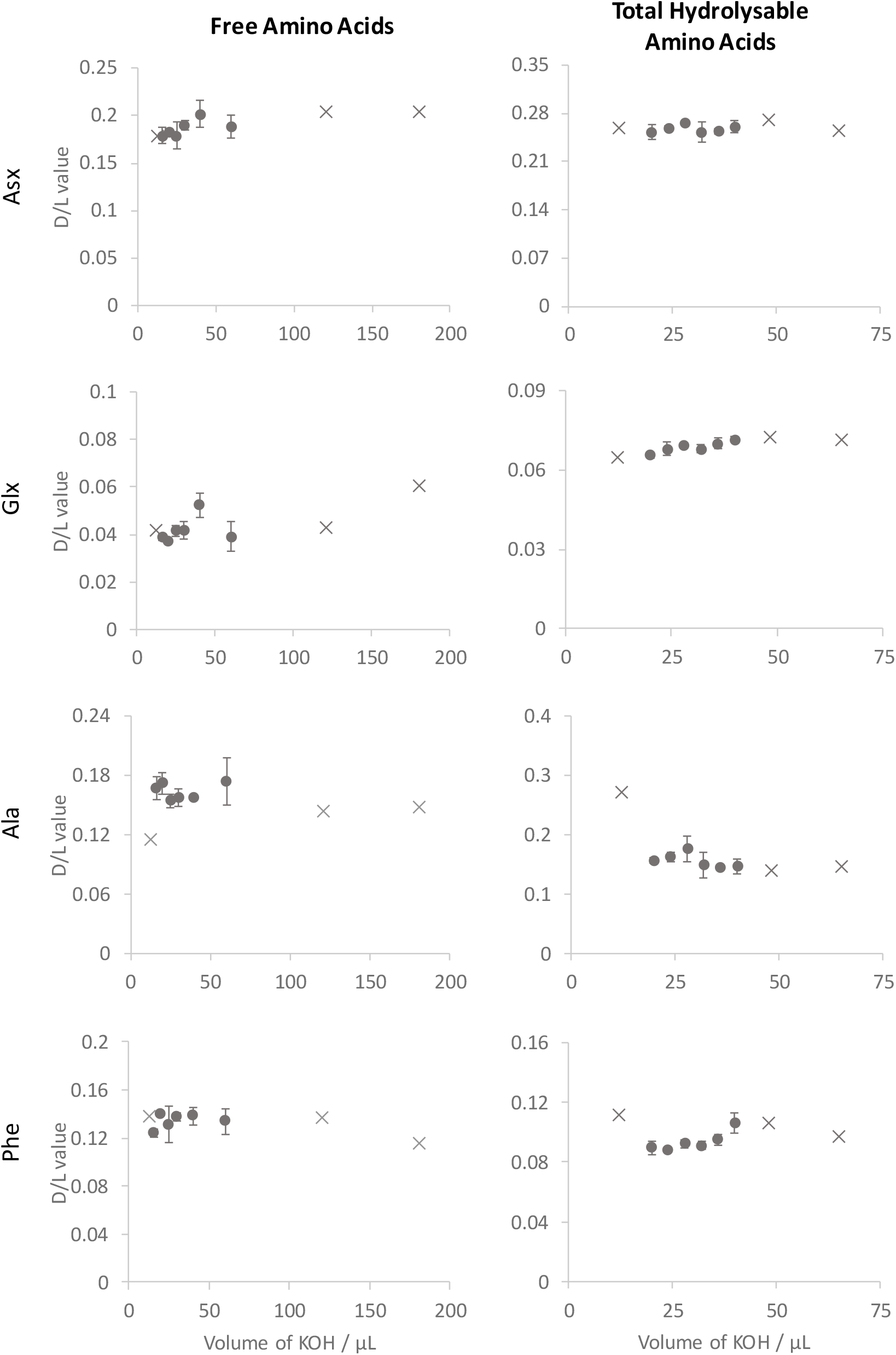
*FAA (Left) and THAA (right) biphasic separation impact on D/L value of amino acids when changing the volume of KOH added). Error bars depict one standard deviation about the mean. Samples that were cloudy prior to HPLC injection have been plotted as crosses, only one sample of each of these volumes was run*.

### 3.3 Impact of the biphasic separation on amino acid composition and D/L value

The biphasic separation of amino acids and phosphate species involves the formation of a gel. The structure of the gel most likely contains a network of inorganic phosphate and calcium ions cross-linked with molecules of water (Gash *et al.*, 2001). It is therefore likely that a proportion of the amino acid content in solution will be retained by the gel and subsequently unanalysed. As each amino acid is chemically unique, they may be retained by the gel to different extents. To test this, a solution with known amino acid composition but different D:L values (1:1 & 4:1) was added topure hydroxyapatite (principal mineral component of enamel) and the amino acid composition was analysed after the optimised biphasic separation.

The amino acid composition of the hydroxyapatite with amino acid refence solutions is broadly unchanged by the biphasic separation procedure (Glx, Ser, L-Thr, Ala, Tyr are biased by <6 %; Figure 5). However, the percentage contribution of Asx to the total amino acid content is slightly reduced and the contribution of the more apolar amino acids (Leu & Ile) is slightly increased. This indicates that there is a slight preferential retention of the more polar amino acids into the gel complex. This preferential incorporation of some amino acids over others may result in inaccurate amino acid compositional data, but it is consistent between analyses and the magnitude of this is unlikely to impact on the estimation of IcPD in fossil samples.

Quantification of the HPLC peak areas of Leu and Ile have been shown to be vulnerable to co-eluting peaks in samples from calcium carbonate-based biominerals (Kaufman & Manley 1998; Powell *et al.*, 2013; Wehmiller, 2013), and this susceptibility also appears to be true for enamel derived samples (Figure 5; Figure 6). Conclusions drawn from these amino acids therefore require caution.

**Figure 5.**
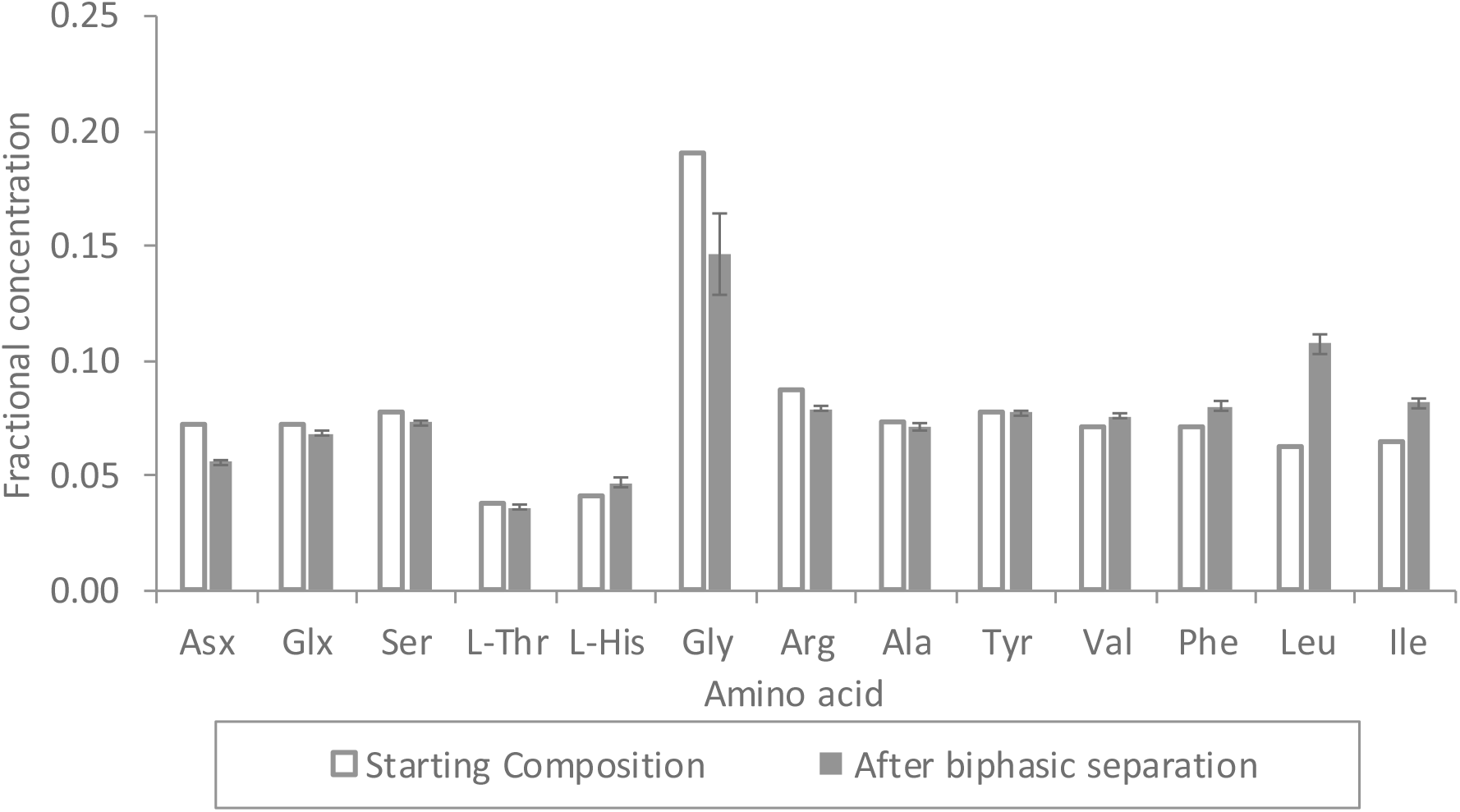
*Fractional concentration of a 1:1 amino acid reference solution with hydroxyapatite that has undergone the biphasic separation of the amino acids from inorganic phosphates (solid bars). The starting composition has been shown for comparison (hollow bars). Error bars depict one standard deviation about the mean. Data was slightly skewed by the higher than expected concentration of Leu*.

#### 3.3.1 Changes in amino acid D/L value induced by biphasic separation

The biphasic separation of amino acids from phosphate species does not alter the D/L value of most amino acids at both low (~0.2) and high (~1) D/L values (Figure 6). This, in conjunction with the IR analysis, indicates that this method of separation is suitable for evaluating amino acids in enamel by RP-HPLC.

**Figure 6.**
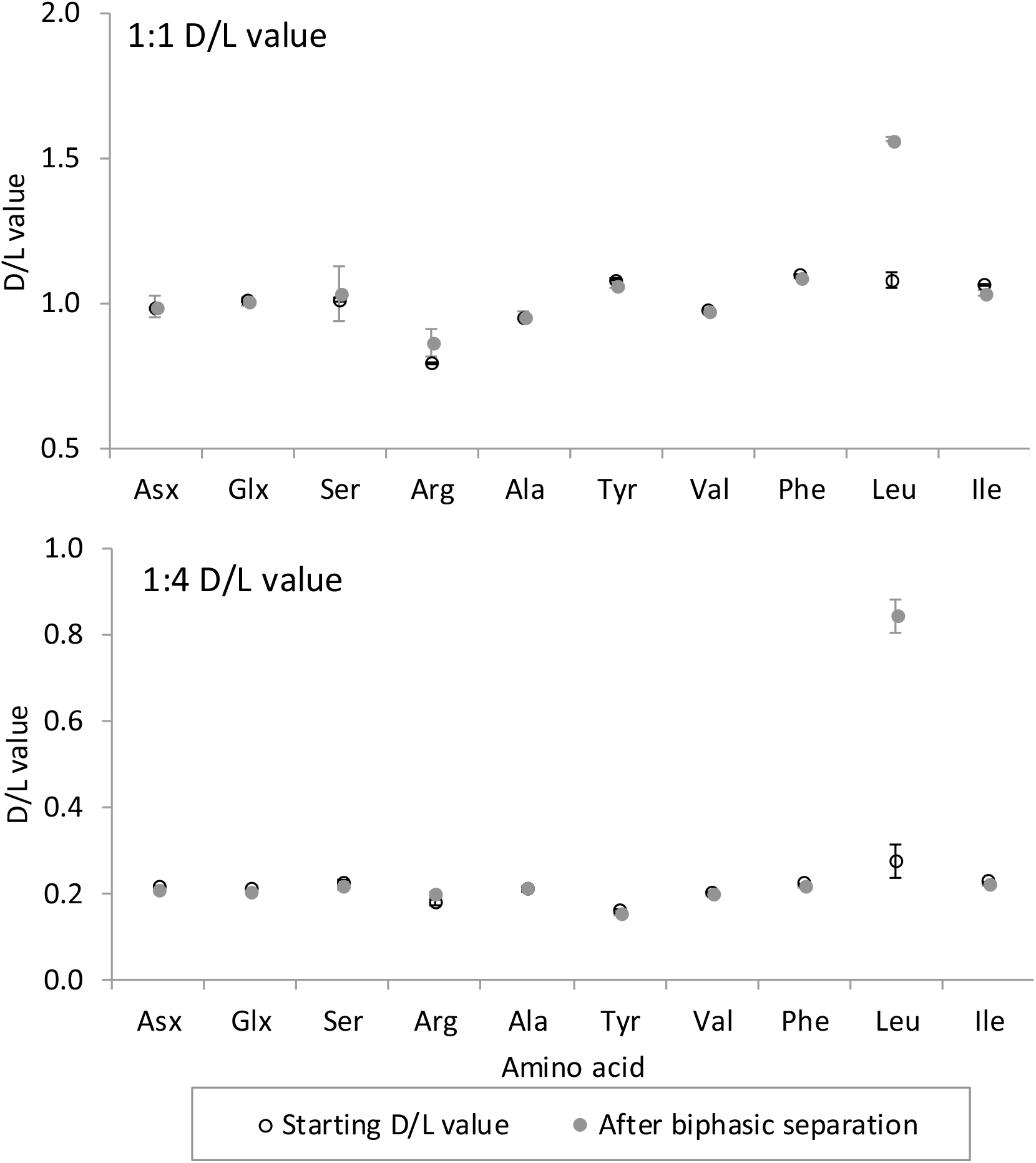
*Amino acid D/L values for hydroxyapatite with amino acid reference solutions. Two different D/L values reference solutions were used to mimic both old (1:1 D/L value; top) and young (1:4 D/L value; bottom) fossil samples. Error bars depict one standard deviation about the mean*.

#### 3.3.2 Estimation of analytical errors

Analytical error here is defined as the variability in amino acid D/L values resulting from the sample preparative procedures and analysis, in this case by RP-HPLC. Most fossil enamel samples can be prepared in duplicate, however, due to the precious nature of some of the samples this may not always be possible. It is essential for geochronological applications that the developed protocol is reproducible. The replicate analysis of eight fossil enamel samples and their standard deviations and coefficient of variance is presented in Table 3. With the exception of Val, the coefficient of variance for most of the amino acids obtained from enamel are on the same order as those values reported for comparably racemized *Bithynia* opercula (Penkman *et al.*, 2013). We therefore conclude that the biphasic separation method developed in this section does enable sufficiently precise amino acid data to be obtained (cf. Table 3 for coefficients of variation). However, it is noted that this degree of precision may not hold true for the entire range of D/L values required for geochronological assessment over Quaternary time scales, so additional analysis of younger and older enamel material would be valuable. Due to the low degree of precision of Val and Tyr D/L values, these amino acids have not been used to build a geochronology.

**Table 3.** *Experimental variability for each amino acid of the optimised enamel AAR protocol on a M. primigenius tooth from the Tattershall Thorpe sand and gravel.*

|  | THAA |  |  | FAA |  |  |
| --- | --- | --- | --- | --- | --- | --- |
|  | Mean D/L |  | Coefficient<br>of variance | Mean D/L |  | Coefficient<br>of variance |
| | value | $\sigma$ | | value | $\sigma$ | |
| Asx | 0.248 | 0.008 | 3.2 % | 0.186 | 0.003 | 1.8 % |
| Glx | 0.070 | 0.003 | 4.6 % | 0.039 | 0.001 | 2.7 % |
| Ser | 0.230 | 0.017 | 7.2 % | 0.413 | 0.020 | 4.9 % |
| Arg | 0.225 | 0.027 | 12.1 % | 0.402 | 0.031 | 7.7 % |
| Ala | 0.154 | 0.009 | 5.8 % | 0.163 | 0.006 | 3.7 % |
| Tyr | - | - | - | 0.124 | 0.051 | 40.7 % |
| Val | 0.134 | 0.027 | 20.2 % | - | - | - |
| Phe | 0.090 | 0.005 | 5.9 % | 0.169 | 0.011 | 6.6 % |
| Leu | - | - | - | - | - | - |
| Ile | - | - | - | - | - | - |

## 4 Isolation of the intra-crystalline fraction

### 4.1 Prolonged oxidative treatment

Previous studies have reported that prolonged oxidative treatment of calcium carbonate based biominerals greatly reduces the amino acid content but leaves a small fraction that is chemically inaccessible (Crenshaw, 1972; Brooks *et al.*, 1990; Sykes *et al.*, 1995; Penkman *et al.*, 2008; Crisp *et al.*, 2013). NaOCl is predicted to oxidise all amino acids and proteins it encounters but will be ineffective on material that is inaccessible. Oxidative pre-treatment of mollusc shell has been shown to improve the accuracy and precision of AAR data, which is vitally important for its application as a tool for geochronology (Penkman *et al.*, 2008; Demarchi *et al.*, 2013a; Ortiz *et al.*, 2018). Powdered enamel was therefore exposed to bleach for increasing lengths of time to test if a similar fraction can be isolated from enamel, and if so, to calculate the optimum length of time to isolate this closed system material.

### 4.2 Optimising the oxidation procedure

#### 4.2.1 Optimisation of the oxidation method

Powdered enamel (~30 mg) from Tattershall Thorpe was weighed into sterile 2 mL microcentrifuge tubes (Eppendorf), and samples were submerged in NaOCl (12 %, 50 µL mg^−1^) for 0, 6, 15, 25, 48, 72 and 238 h. The NaOCl was removed and the powdered enamel washed five times with HPLC-grade water. A final wash with methanol was performed before the samples were left to air dry overnight. Samples were then purified using the optimised biphasic procedure (section 2.4.2) and analysed by RP-HPLC (Section 2.5). Samples were prepared with 4 replicates.

#### 4.2.2 Changes in amino acid concentration with prolonged oxidative treatment

The FAA concentration remains constant up to 6 h, increases by 15 h, before returning to approximately the initial concentrations (Figure 7). The concentration of the FAA does not decrease when exposed to NaOCl which indicates that the FAAs are inaccessible to the NaOCl and potentially retained in a closed system. The enamel used in this set of experiments was ~191 ka in age, so it is likely that most of the open system FAAs would have leached out whilst in the depositional environment, thus leaving a mostly closed system fraction of FAA resistant to oxidative treatment.

**Figure 7.**
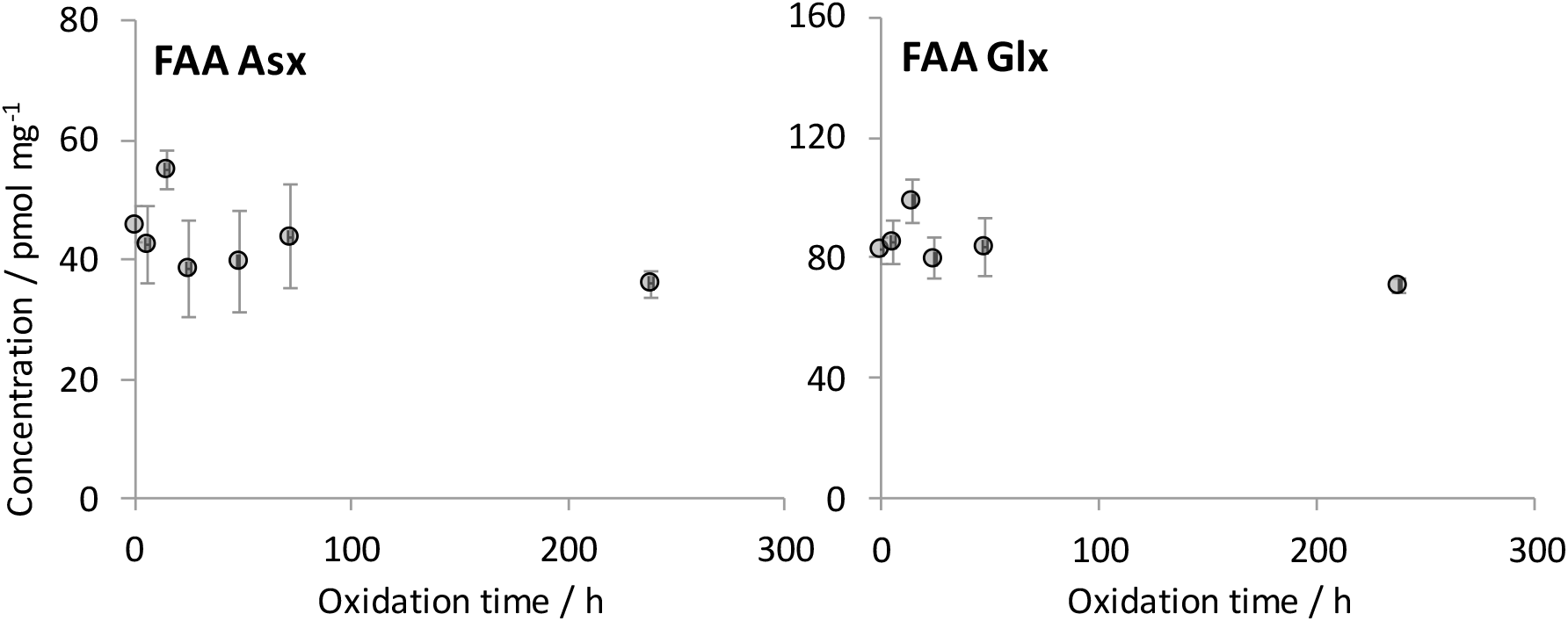
*FAA concentrations from powdered *M. primigenius* enamel of Asx (left) and Glx (right) after prolonged exposure to a strong oxidant (NaOCl). Error bars depict one standard deviation about the mean (n=4 or 5)*.

The concentration of THAA in enamel decreased rapidly by ~25 % during the initial 6 h of exposure to NaOCl (Figure 8). This rapid loss of organic matter has been reported for other biominerals (Bright & Kaufman, 2011; Hendy *et al.*, 2012; Crisp *et al.*, 2013) and has been attributed to the oxidation of easily accessible organic matter, believed to be present in an open system found between crystallites (Towe and Thomson, 1972). After this point a more gradual decrease in concentration was observed; it is likely that a fraction of organic matter present between crystallites is resistant to the initial exposure of strong oxidant and therefore enamel requires further exposure to fully remove this material. After 72 h of exposure to bleach there is only minimal additional loss of the THAAs.

**Figure 8.**
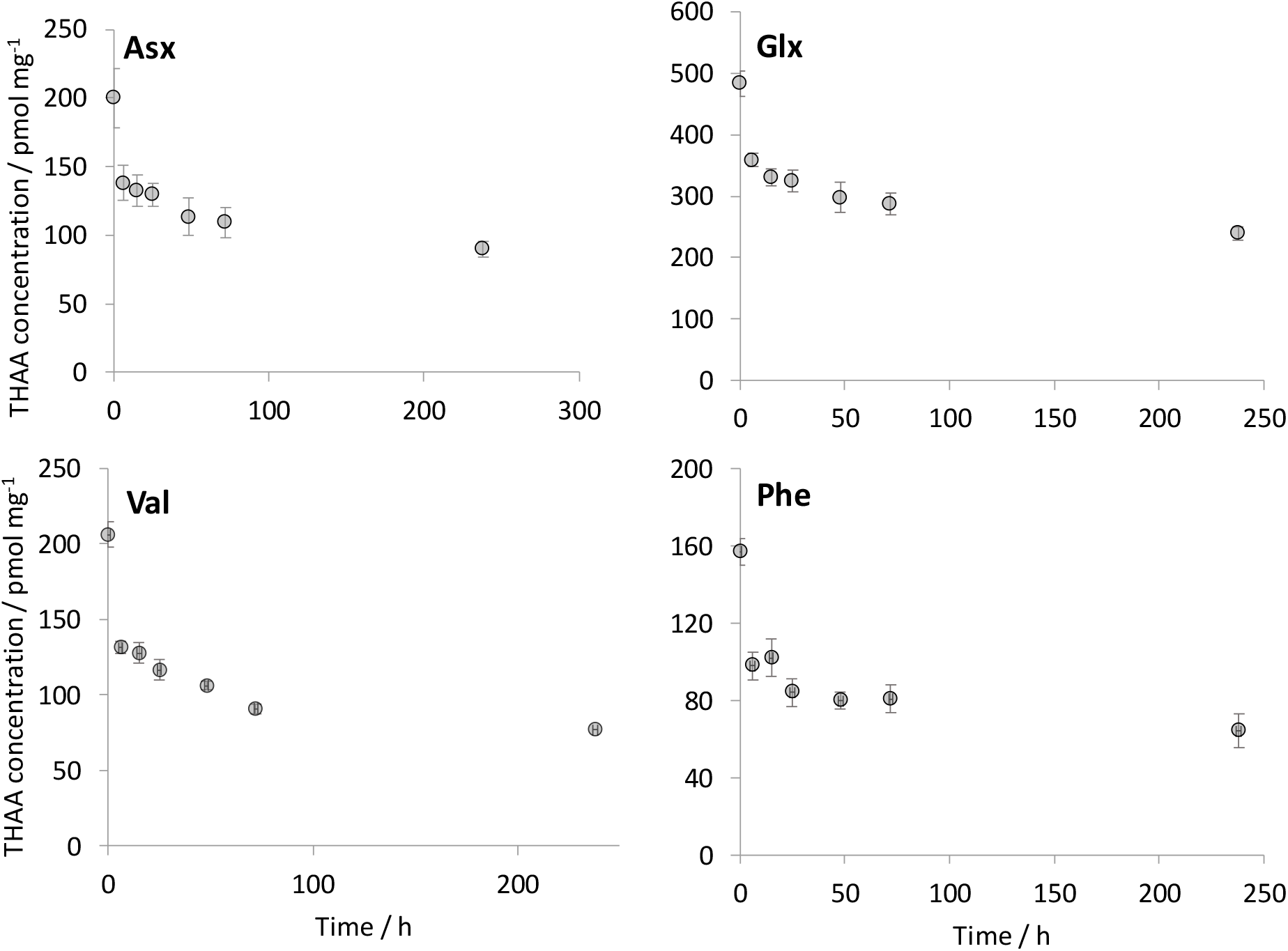
*THAA concentrations from powdered *M. primigenius* enamel of Asx (top left), Glx (top right), Val (bottom left) and Phe (bottom right) after prolonged exposure to a strong oxidant (NaOCl). Error bars depict one standard deviation about the mean (n=4 or 5)*.

#### 4.2.3 Effect of increased oxidation time on D/L values

The FAA D/L values for Asx, Glx, Ala and Phe remain relatively constant with increasing exposure time (Figure 9), which is consistent with the trends observed for the FAA concentrations (Figure 7) and indicate the FAA fraction is in a stable, inaccessible environment.

The THAA D/L values for some of the amino acids does not change (Glx and Phe), indicating that the organic material removed by the prolonged oxidation is at a similar level of degradation to that which has been isolated. However, this will most likely not always be the case (e.g. Penkman et al 2007, Ortiz *et al.*, 2018).

The THAA D/L values for Asx decline gradually to 72 h and then only minimal further reduction is observed. The extent of bound Asx racemization is initially greater than for free Asx (Figure 9). The Asx THAA concentration decreases and the FAA concentration is stable with exposure to NaOCl up until 72 h (Figure 7 and 3) and thus NaOCl is removing the more racemised bound Asx, causing the gradual decrease in Asx THAA racemization.

Studies of molluscan shells and ostrich eggshell have reported that after the intra-crystalline fraction has been isolated, a gradual increase in D/L value is observed with increasing exposure time (Penkman *et al.*, 2008; Crisp *et al.*, 2013). In aqueous solutions, bleach forms a weak acid and this increase in D/L value is thought to be caused by gradual etching of the calcium carbonate matrix, inducing racemization. This D/L pattern is not observed in the sample of enamel, but gradual loss of amino acids might be caused by the same slow etching of the mineral matrix.

The extent of Asx and Glx racemization in the FAA fraction is less than in the THAA fraction (Figure 9). This is the opposite of the patterns observed in other biominerals (e.g. Penkman *et al.*, 2008; Tomiak *et al.*, 2013) and indicates that the bound amino acids are more racemised than the FAAs. FAAs are expected to be more racemised than THAAs, because with the exceptions of Asx and Ser, most amino acids are unable to racemise in-chain (Stephenson & Clarke, 1989; Takahashi, *et al.*, 2010; Demarchi *et al.*, 2013b). Amino acids situated at the terminal position of peptides are thought to have the lowest activation of racemization and thus racemization occurs rapidly (Kriausakul & Mitterer, 1978; Stephenson & Clarke, 1989). Free amino acids form as a result of hydrolysis of these more highly racemised amino acids and thus are expected to have a greater extent of racemization. In enamel, in-chain racemization may account for the reversal of trend for Asx but not for Glx, so this is a very interesting observation that may indicate enamel peptides are breaking down in a different way to other biominerals, possibly due to the maturation process *in vivo* (Robinson *et al.*, 1995; 1998).

**Figure 9.**
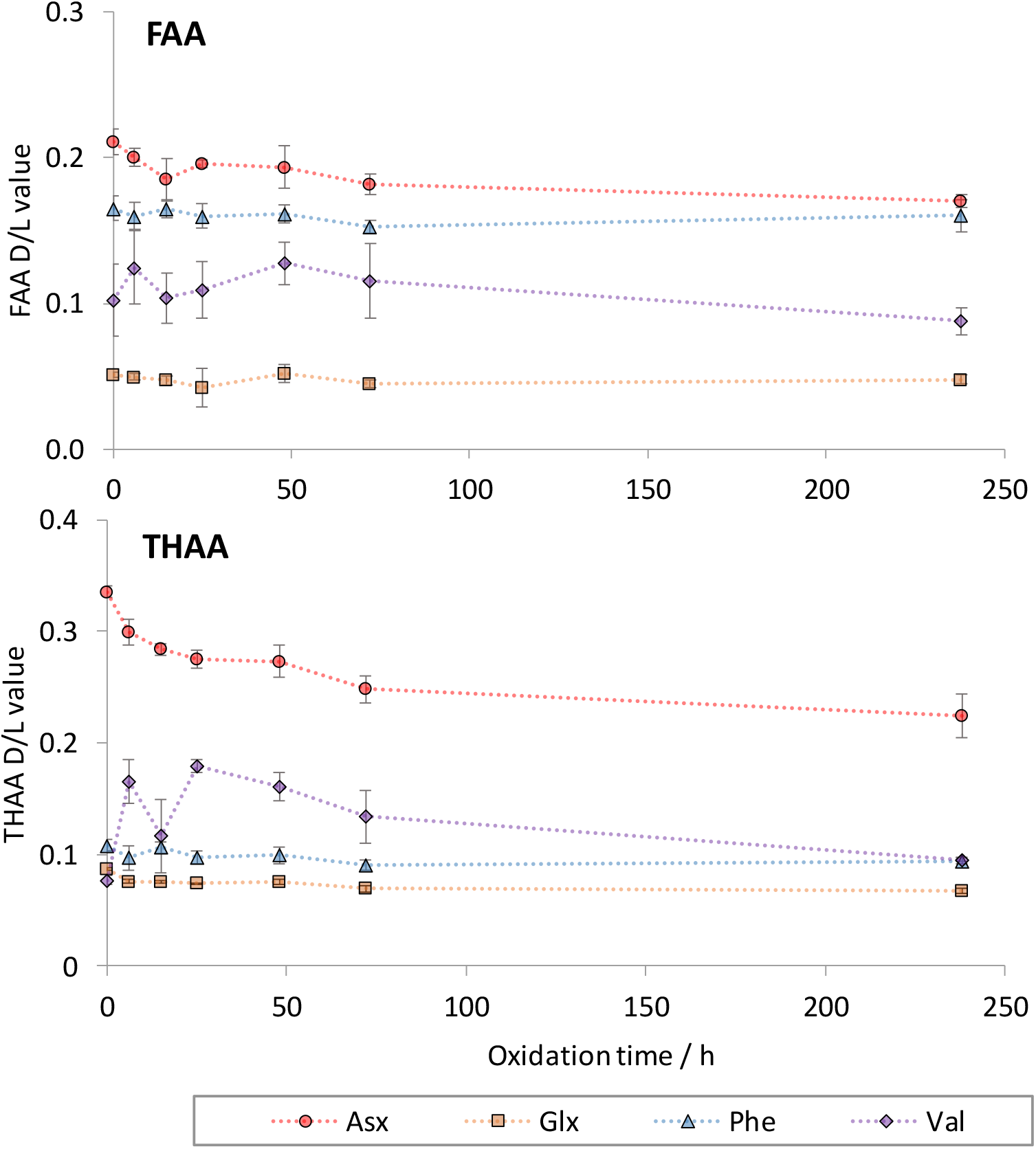
*Effects of oxidative treatment on the extent of racemization in powdered *M. primigenius* enamel for FAA (Top) and THAA (bottom) fractions. Error bars depict one standard deviation about the mean (n= 4 or 5)*.

#### 4.2.4 Intra-crystalline enamel composition

The composition and role of protein in immature dental enamel have been widely studied (Margolis *et al.*, 2006), but little is known about the proteinaceous material that can be recovered from mature/fossil enamel. Mature enamel is acellular and contains a complex mixture of peptides, amino acids and proteins originating from a variety of different dental developmental processes (Robinson *et al.*, 1995). Collagen contamination from the surrounding dentine and cementum poses a risk to enamel AAR dating. Collagen has a characteristically high proportion of Gly (~33%), proline (~12%) and hydroxyproline (10 %; Poinar & Stankiewicz, 1999). The method of analysis used in this study cannot detect proline or hydroxyproline, but the relatively low levels of Gly (< 33 %) observed in the bleached THAA fraction of mammoth enamel (Figure 10) indicates that the amino acids analysed by this procedure have likely originated from mature enamel proteins rather than collagen. The enamel profile also matches that of previously reported fossil enamel from a closely related taxon, *Mammut sp*. (mastodon; Figure 10).

**Figure 10.**
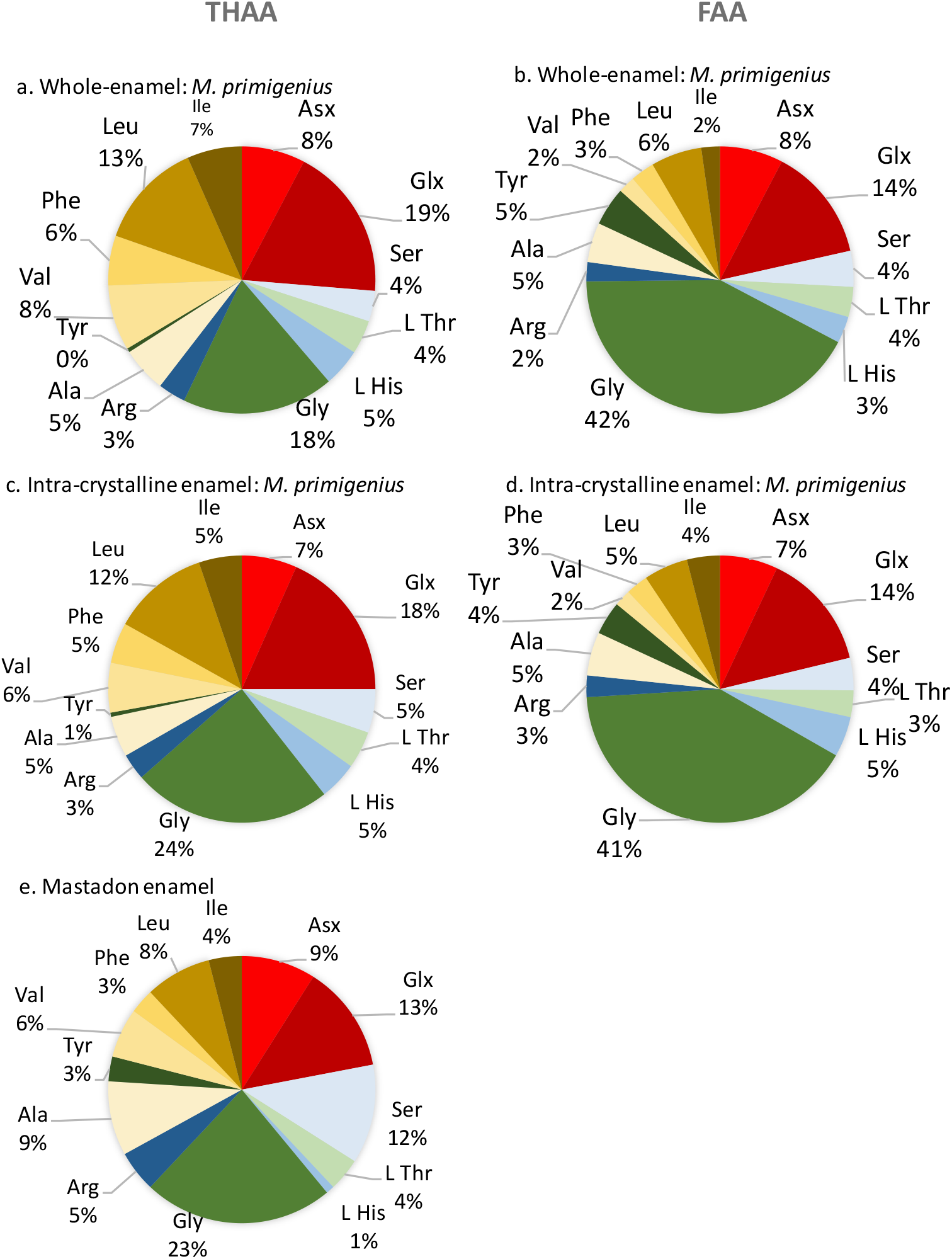
*FAA (Right) and THAA (left) composition of *M. primigenius* enamel from the Thorpe sand and gravel at Tattershall, from both whole-enamel (top), intra-crystalline (middle), and previously reported mastodon enamel (bottom; Doberenz *et al.*, 1969). Amino acids are coloured differently based on their side-chain properties: Ionisable side-chains with a negative charge (red), ionisable side-chains with a positive charge (blue), non-ionisable side-chains that are polar (yellow) and non-ionisable side-chains that are non-polar (green)*

The THAA composition of the unbleached enamel is similar to that of the bleached (intra-crystalline; Figure 10). This suggests that they are made up of an analogous set of proteins. However, there is an increase in the relative contribution of Gly in the intra-crystalline fraction. The concentration of the Gly rich FAA fraction has been shown to be stable upon increased exposure time to bleach (Figure 9), whilst the THAA concentration decreases. The THAA content is comprised of both the free and bound amino acid fractions, so the change in the THAA composition is likely to be due to the loss of the more Gly-poor bound amino acids upon bleaching.

The similar compositional change of amino acids during bleaching has been observed in ostrich eggshell (Crisp *et al.*, 2013) but not for other biominerals (Ingalls et al, 2003; Penkman *et al.*, 2008; Demarchi *et al.*, 2013c). It has been observed that acid-rich proteins preferentially bind to calcite, enhancing their contribution to the intra-crystalline fraction (Marin *et al.*, 2008). However, this has not been observed for enamel, and is potentially a result of different pathways for the structural formation of enamel or for the degradation of enamel proteins.

## 5 Elevated temperature experiments and fossil analyses to test for closed system behaviour

The process of racemization at ambient diagenetic temperatures occurs over geological timescales, in some cases taking millions of years to equilibrate (Hare & Mitterer, 1967). To study the reaction kinetics of IcPD/racemization in a laboratory, this process needs to be accelerated and all the amino acids in the system need to be accounted for (Collins & Riley, 2000). Bleached and unbleached samples have been analysed to compare the behaviour of the inter- and intra-crystalline fractions of amino acids in enamel (section 4). However, bleaching treatment may not isolate a purely intra-fraction as bleached biominerals may also contain quantities of highly resilient proteins from the inter-crystalline fraction (Bright & Kaufman, 2011).

Previous studies of the long-term diagenesis of proteins in other biominerals (e.g. mollusc shells, Penkman *et al.*, 2008; OES, Crisp *et al.*, 2013; corals, Tomiak *et al.*, 2012) have heated samples isothermally in sealed vials containing water; analysis of both the biomineral and the supernatant water thus accounts for all amino acid degradation products. Consideration of the reaction kinetics of the proteins and amino acids in enamel under laboratory conditions can help inform our understanding of some of the trends observed in fossil data. High temperature kinetic and leaching experiments have therefore been undertaken on both modern and fossil enamel to examine their degradation behaviours.

### 5.1 Methods for testing for a closed system

Approximately 10 mg of bleached and unbleached powdered tooth enamel from both *M. primigenius* from the North Sea and a modern *E. maximus* was weighed into sterile glass ampoules. 500 μL of HPLC grade water was added and the glass vials were sealed. The ampoules were placed in an oven and heated isothermally at 80, 110 or 140 °C for varying times (Table 4). Triplicate samples were analysed for each time-point.

**Table 4.** *Conditions for the heating of powdered enamel. Triplicate samples were prepared for each time point. Bleached samples were agitated in NaOCl for 72 h according to section 4.2.1.*

| Sample | Bleached | T / °C | Heating time / h |  |  |  |  |  |  |  |  |  |  |  |
| --- | --- | --- | --- | --- | --- | --- | --- | --- | --- | --- | --- | --- | --- | --- |
| Modern Elephant | Yes | 140 | 0 | 1 | 2 | 4 | 6 | 8 | 24 | 48 | 72 | 96 | 120 |  |
| Modern Elephant | No | 140 | 0 | 1 | 2 | 4 | 48 |  |  |  |  |  |  |  |
| Modern Elephant | Yes | 110 | 0 | 24 | 48 | 148 | 240 | 384 | 480 | 720 | 840 | 963 | 1200 | 2688 |
| Modern Elephant | Yes | 80 | 0 | 24 | 96 | 148 | 480 | 720 | 963 | 1442 | 2160 | 3600 | 4560 | 5760 |
| North Sea | Yes | 140 | 0 | 1 | 4 | 6 | 8 | 24 | 48 |  |  |  |  |  |
| North Sea | No | 140 | 0 | 1 | 6 | 48 |  |  |  |  |  |  |  |  |

After the allotted time, the glass ampoules were unsealed and the supernatant water split into two aliquots (100 μL). One fraction was analysed for the free amino acids present in the water (FAAw) and the other was analysed for the total hydrolysable amino acid content in the water (THAAw). To THAAw samples, HCl (6 M, 20 μL per maximum theoretical mg equivalent of enamel) was added and placed in an oven for 24 h at 110 °C in N_2_ purged vials. To remove the HCl, samples were placed in a centrifugal evaporator. FAAw samples were dried in a centrifugal evaporator with no further sample preparation.

Powder samples were weighed out into two fractions for the analysis of the free and total hydrolysable amino acid content in the powders (FAAp and THAAp respectively) according to the optimised procedure outlined in sections 2.4.1–2.5. All sub-samples were analysed by RP-HPLC (section 2.5).

The supernatant water of bleached modern elephant enamel powder isothermally heated at 140 °C for 48 h was concentrated 10-fold and the resultant residue imaged by transmission electron microscopy (TEM). Formvar/Carbon 200 mesh Copper grids were plasma cleaned for 10 seconds in a Harrick Plasma PDC-32G-2. A 3 μL volume of sample was applied to the grid and either removed by blotting after 1 min and left to air dry, or left to dry for 30 min without blotting to increase the sample deposited on the grid. Grids were imaged using a Tecnai G_2_ 12 BioTWIN microscope operating at 120 kV with a Tungsten filament. Some of the TEM plates were overloaded with crystals and thus damaged.

#### 5.1.1 Fossil enamel pilot study

Enamel from: a modern elephant and mammoths from the North Sea, Balderton, Tattershall Thorpe, Crayford, Ilford, Swanscombe, Sidestrand and the Norwich Crag formation were individually prepared and analysed for their intra-crystalline amino acid content by RP-HPLC according to the optimised protocols outlined in sections 2.2–2.5.

### 5.2 Intra-crystalline vs whole-enamel D/L values

As expected, in modern elephant enamel, THAA D/L values for intra-crystalline Asx, Glx, Ala and Phe increase over time when heated isothermally at 140 °C (Figure 11). However, the D/L values for the modern elephant whole-enamel (unbleached) proteins initially increased and then plateaued, deviating greatly from the trend observed for the intra-crystalline fraction. Comparable experiments of other biominerals that simulate protein degradation have shown similar patterns, which have been attributed to the early leaching of the more highly racemised FAAs from the whole-shell fraction of gastropod, and ostrich eggshell (Crisp *et al.*, 2013; Ortiz *et al.*, 2018). In contrast, this process does not occur in the intra-crystalline fraction because it exhibits closed system behaviour. The precision of the intra-crystalline D/L values are higher than for the whole-enamel data (cf. error bars, Figure 11), with the higher variability for the whole-enamel data likely to be a consequence of differential leaching of open system material.

**Figure 11.**
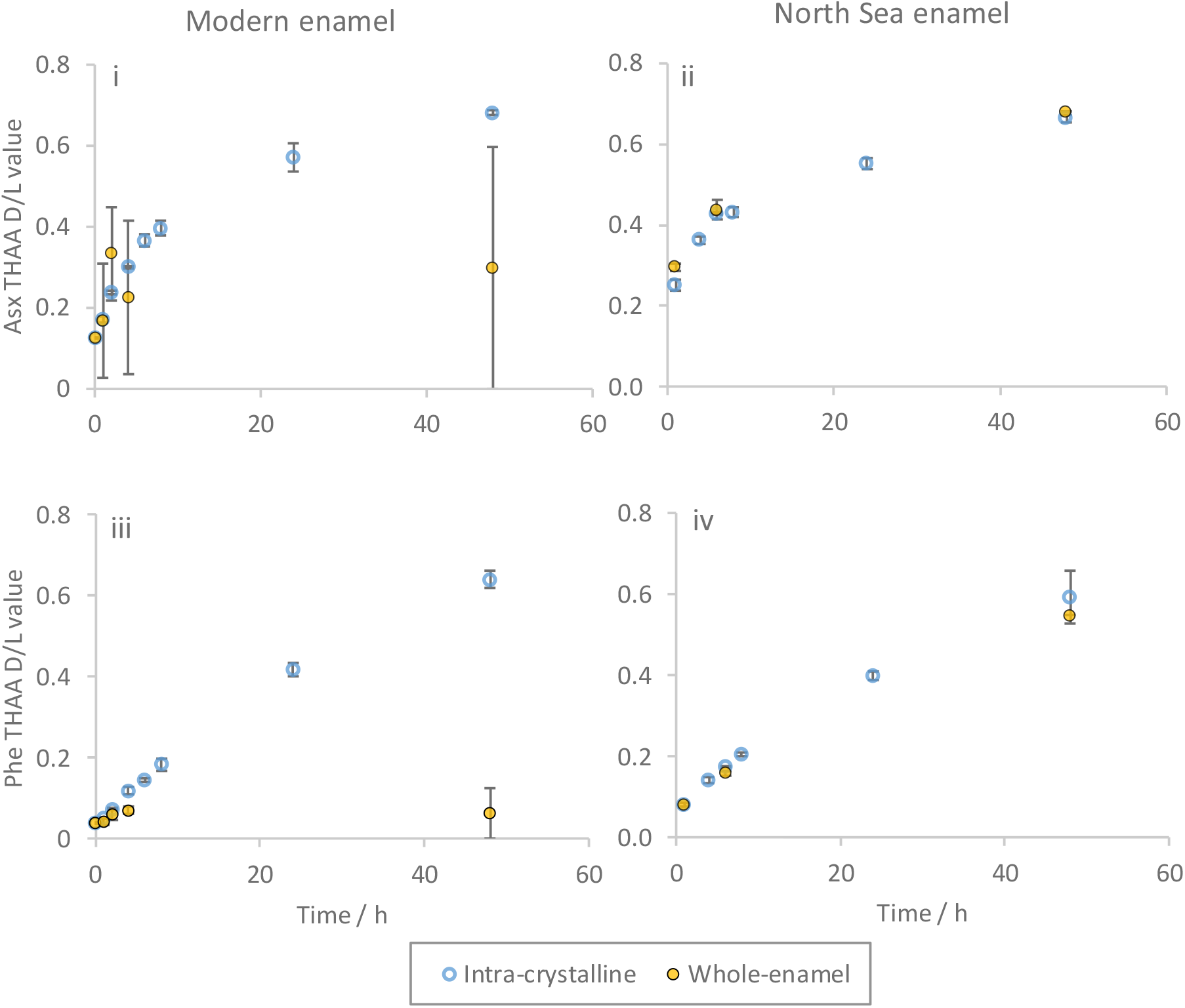
THAA D/L value of Asx (top) and Phe (bottom) for bleached (intra-crystalline) and unbleached (whole-enamel) modern *E. maximus* enamel (i and ii) and North Sea *M. primigenius* enamel (ii and iv) with increasing time when isothermally heated at 140 °C. Error bars depict one standard deviation about the mean (Modern enamel n = 2-6, North Sea enamel n = 2-3).

In contrast to the modern enamel, there is no significant difference in extent of THAA racemization between bleached and unbleached North Sea enamel heated at 140 °C (Figure 11). It is likely that the fossil enamel has already lost most of the leachable organics and thus the whole-enamel protein composition is likely to be instead dominated by the intra-crystalline amino acids and thus the bleached and unbleached fractions are behaving in a similar manner.

### 5.3 Leaching of amino acids into the supernatant water

It is expected that during an accelerated degradation experiment, open system amino acids will leach out of the enamel powder; this diffusive loss would also be relevant in the depositional environment (Collins & Riley, 2000). Previous studies have shown that bleaching can significantly reduce the quantity of leached amino acids in similar IcPD experiments (Penkman *et al.*, 2008; Crisp *et al.*, 2013; Ortiz *et al.*, 2018). To quantify the extent of leaching, the enamel powder was heated in water so that leached material could be recovered through subsequent analysis of the water (section 5.3.1).

#### 5.3.1 Concentration of amino acids in the supernatant water

The concentration of leached amino acids in the supernatant water of bleached enamel is notably lower than observed in the waters of unbleached enamel (Figure 12 & 14). This implies that most of the amino acids removed by bleaching are open system and thus can leach out of the enamel during these degradation experiments (Penkman *et al.*, 2008). This is supported by the trend in D/L values observed for the powdered modern elephant enamel (Figure 11), which suggest the unbleached enamel is losing the more highly racemised amino acids during these experiments.

**Figure 12.**
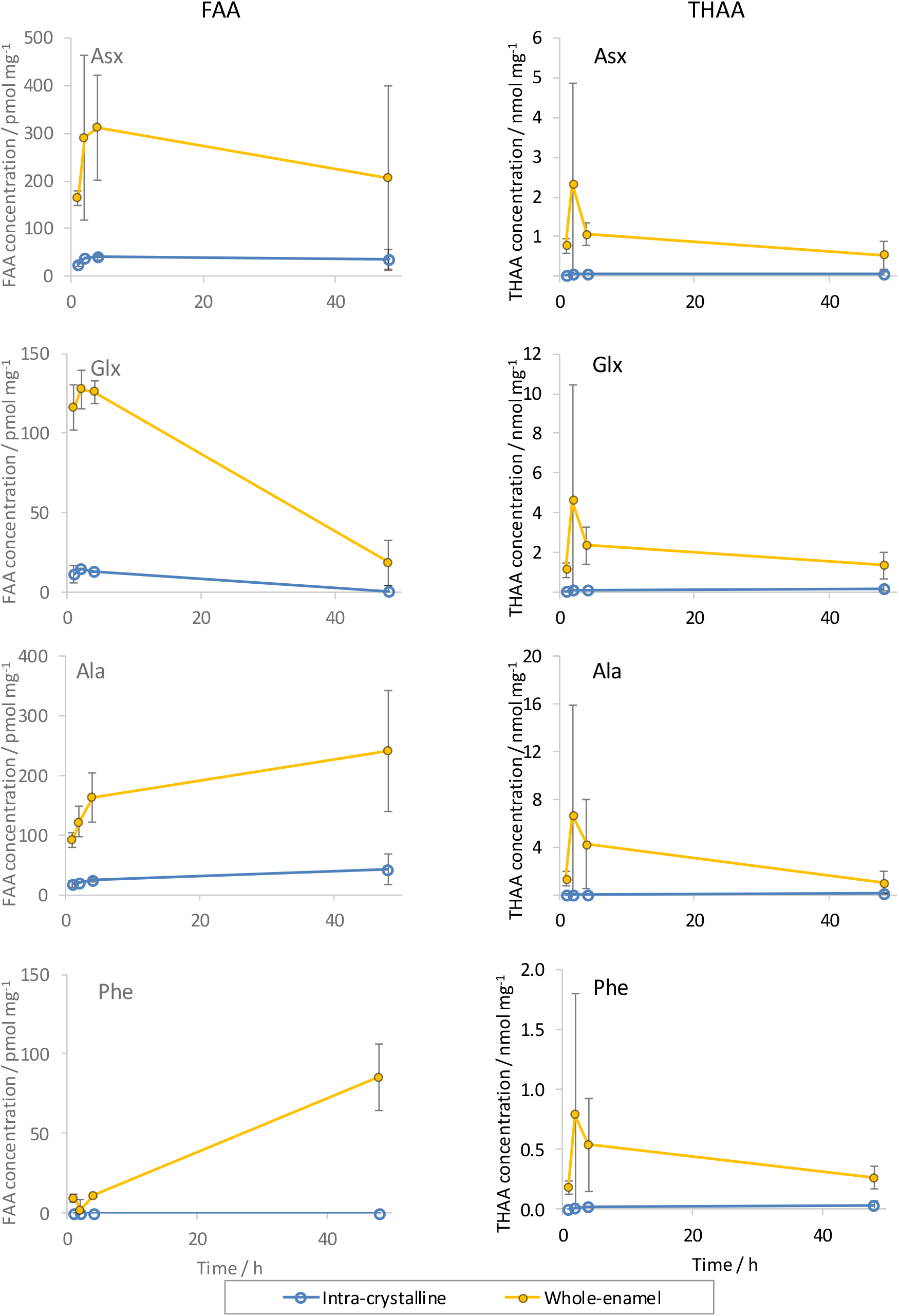
*Concentration of FAA (left) and THAA (right) in supernatant water from heating experiments conducted at 140 °C for both bleached (blue) and unbleached (orange) modern elephant enamel. Error bars show one standard deviation about the mean (n = 2-6).*

Higher concentrations of amino acids were also present in the supernatant waters of unbleached mammoth enamel from the North Sea than in those of bleached enamel (Figure 13), which agrees with the trends observed for modern elephant enamel (Figure 12). Low concentrations of amino acids were also detectable in the supernatant waters from the bleached modern and North Sea enamel. Mean percentage concentration of Asx, Glx, Ser, Ala, Arg, Ala Tyr, Val & Phe in the supernatant waters of enamel after 72 h were ~5 % of the total starting concentration, which is similar to those recorded for gastropod *Phorcus lineatus* (ca. 5%; Ortiz *et al.*, 2018).

**Figure 13.**
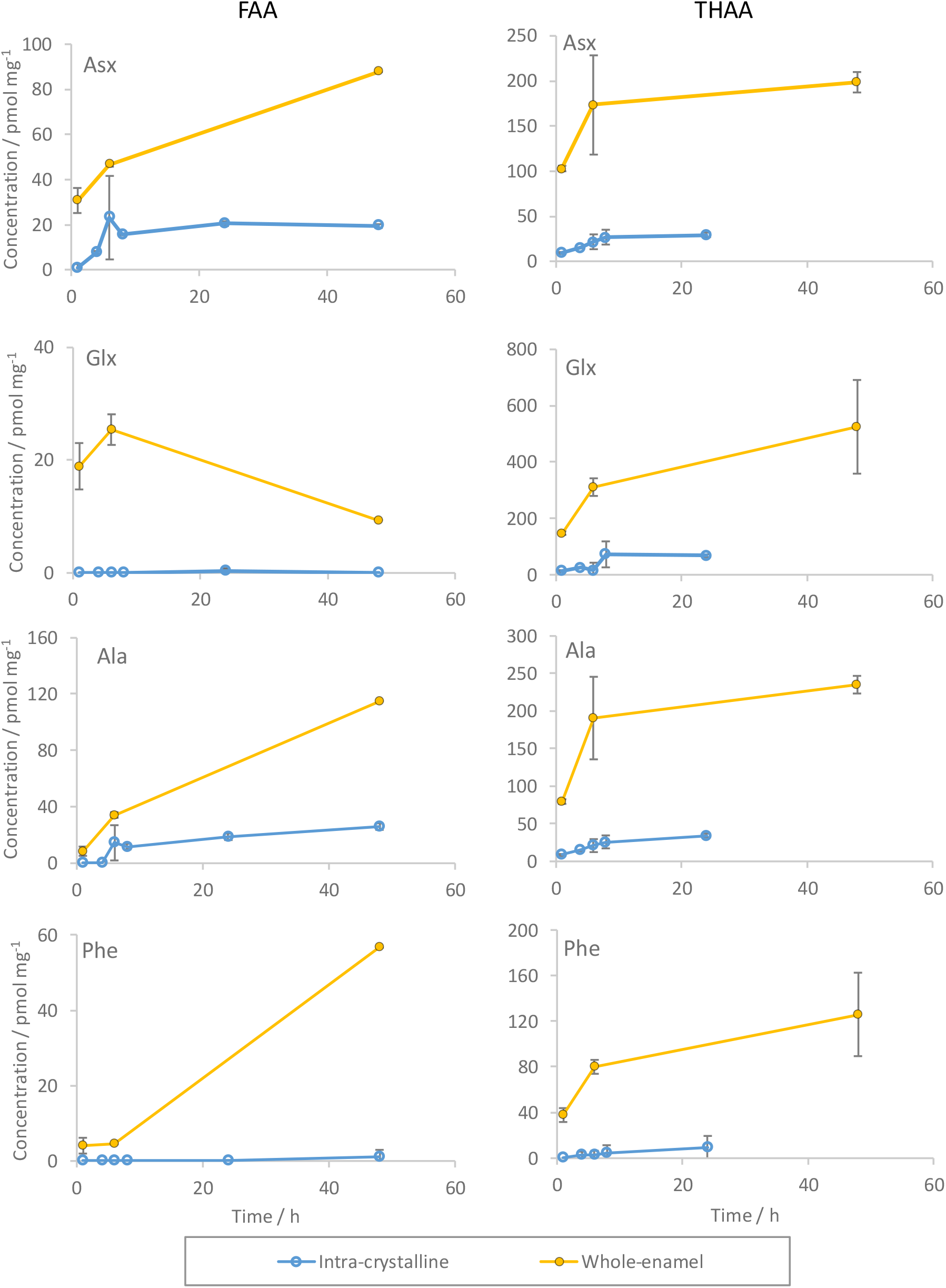
*Concentration of FAA (left) and THAA (right) in supernatant waters of heating experiments conducted at 140 °C for both bleached (Blue) and unbleached (orange) North Sea *M. primigenius*. Error bars depict one standard deviation about the mean (n-2-3).*

However, the enamel powder used in these elevated heating experiments is very fine and thus small particles can be readily disturbed into the supernatant waters, thereby potentially accounting for some of the amino acids in the water fractions. To combat this, the vials were centrifuged prior to extraction, but this may have not been sufficient. The dried supernatant water of a modern elephant sample was therefore imaged by TEM to investigate if crystalline material was present in the supernatant waters of these experiments.

As the supernatant water was concentrated, the solution became cloudy, indicating the presence of solid material. A large range of crystal sizes from tens of μm to ~10 nm was observed by TEM (Figure 14). The abundance of crystalline artefacts indicates that despite the careful attempt to isolate supernatant water only, powdered enamel is still present in the supernatant waters, and thus will contribute to the total FAAw and THAAw contents. This evidence does not preclude the possibility there are leached amino acids in the supernatant waters, but this observation means that we are limited in differentiating leached amino acids from powder contamination of the supernatant water in this study. The concentrations of amino acids in the supernatant water are therefore upper estimates of leaching and the actual percentages of leached amino acids is likely to be lower. The results of these leaching experiments indicate that bleaching does significantly minimise leaching during high temperature diagenesis in both fossil and modern material.

**Figure 14.**
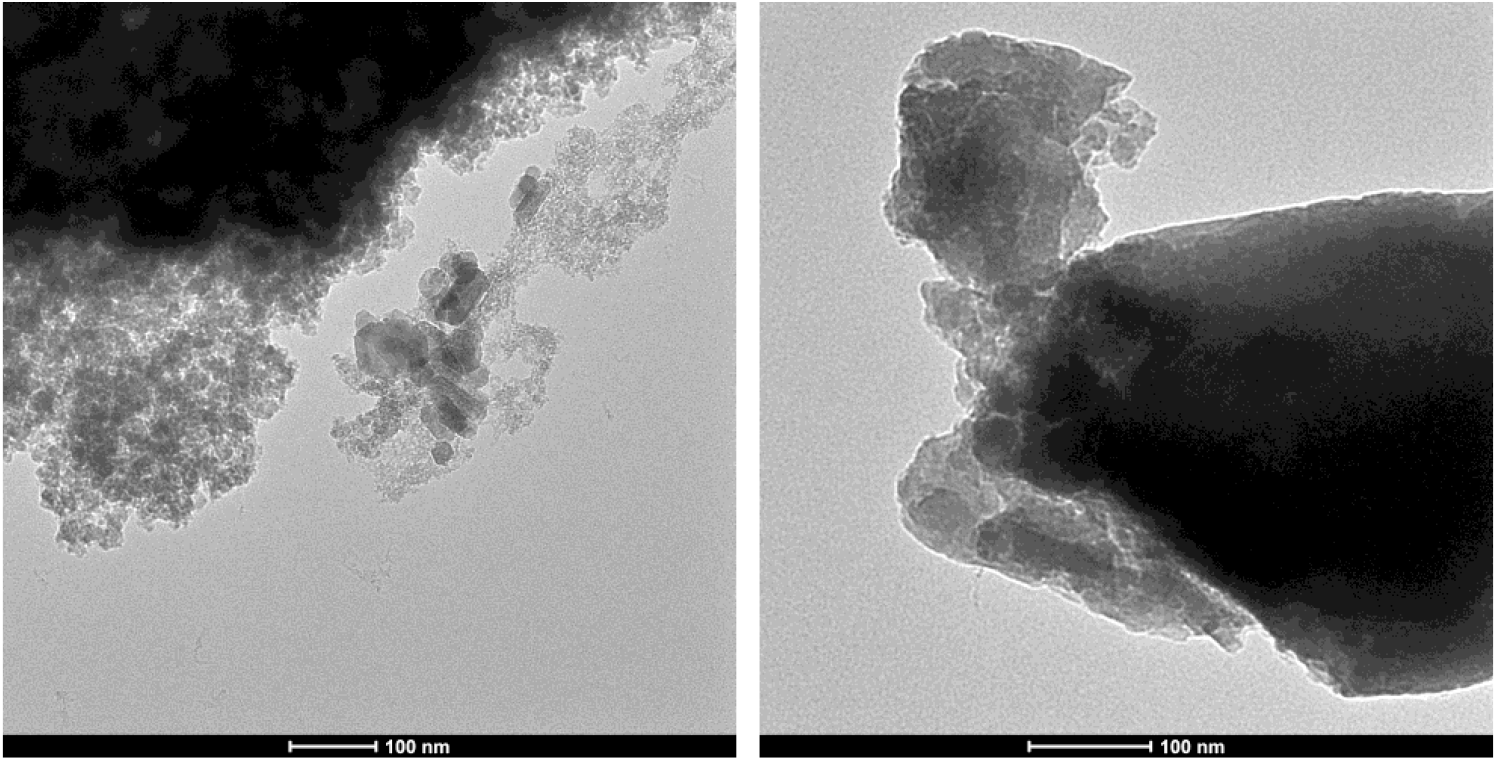
*TEM images of inorganic crystals in the supernatant water of modern elephant enamel powder isothermally heated at 140 °C for 48 h. Supernatant water was left to dry for 30 min on the copper grid.*

### 5.4 Elevated temperature FAA vs THAA D/L

In a closed system, all the diagenetic products of protein degradation are retained within the biomineral up until analysis (Collins & Riley, 2000), the implication of which is that FAA and THAA D/L values should be highly correlated, if enamel is a closed system.

The FAA and THAA D/L values of Asx, Glx, Ala and Phe are highly correlated at all the temperatures studied (Figure 15). The D/L values for some of the least racemised FAAs could not be determined due to very low concentrations of the D isomer and have therefore been plotted along the axis. The high degree of correlation indicates that the FAA are retained by the biomineral and that leaching is not occurring. This supports the assertion that the enamel proteins isolated via bleaching are effectively operating as a closed system.

In calcium carbonate-based biominerals, the extent of FAA racemization is greater than for the THAA, because most amino acids (except Asx and Ser; Stephenson & Clarke, 1989; Demarchi *et al.*, 2013b) are unable to racemise when bound in chain (Mitterer & Kriausakul, 1984). Except for Phe, this trend has generally not been observed to be true for enamel during these degradation experiments (Figure 15). The extent of racemisation observed for the FAA and THAA fractions of Glx and Ala are very similar and increase in a roughly 1:1 ratio (Figure 15). Mature enamel contains shorter chain peptides than most biominerals due to its maturation processes (Robinson *et al.*, 1995), so the lower average chain length peptides in enamel might explain the similar rates of racemisation between the THAA and FAA fractions. The Asx THAA D/L values are greater than the corresponding FAA values (Figure 15), likely to be due to in-chain racemization (Stephenson & Clarke, 1989).

**Figure 15.**
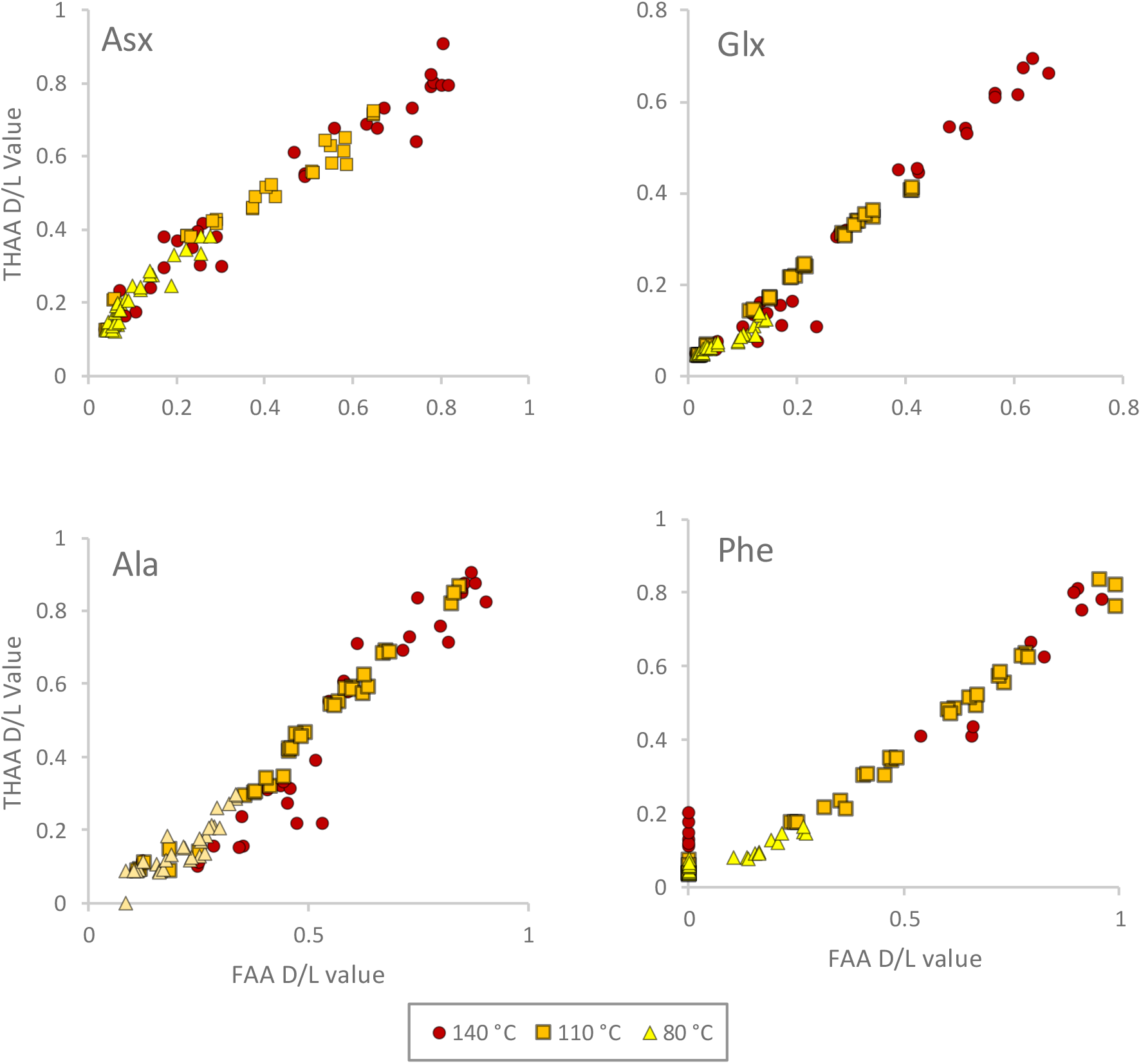
*FAA D/L vs THAA D/L values for bleached enamel in 140 °C, 110 °C and 80 °C heating experiments for Asx (top left), Glx (top right), Ala (bottom left) and Phe (bottom right). Points plotted on the y-axis are due to concentration of the D isomer being below the limits of detection.*

### 5.5 Intra-crystalline enamel amino acids: kinetic behaviour

Except for Ser, a predictable increase in the extent of racemization was observed in all the amino acids studied (Asx, Glx, Ala, Val and Phe) across the range of temperatures (80, 110 and 140 °C; Figure 16). The racemization of Ser increases rapidly and then falls to lower levels of racemization, which is linked to in-chain racemization and chemical instability of Ser respectively (Akiyama, 1980; Takahashi *et al.*, 2010). This trend of Ser racemization in the intra-crystalline fraction has been observed in elevated temperature studies of other biominerals (Kaufman, 2000; Penkman, *et al.*, 2008; Demarchi, *et al.*, 2013c; Orem & Kaufman, 2011) and is supported by observations in the fossil record (Preece & Penkman, 2005).

The mechanism for the racemization of FAAs is thought to predominantly proceed by a carbanion intermediate. The electron-withdrawing and resonance stabilisation capabilities of the substituents attached to the α-carbon are the principal factors in determining the relative order of rates of racemization (Shou & Bada, 1980). However, in a biomineral, the complexity of peptide-bound amino acid diagenesis means that the primary sequence and the chemical environment can also influence relative rates of racemization (Collins *et al.*, 1999). The observed relative order of the rates of racemization of FAAs in solution has been reported to be Asp > Phe > Ala > Glu > Val (Smith & Evans, 1980); the heated experiments conducted on enamel in this study show broadly similar trends (Figure 16).

**Figure 16.**
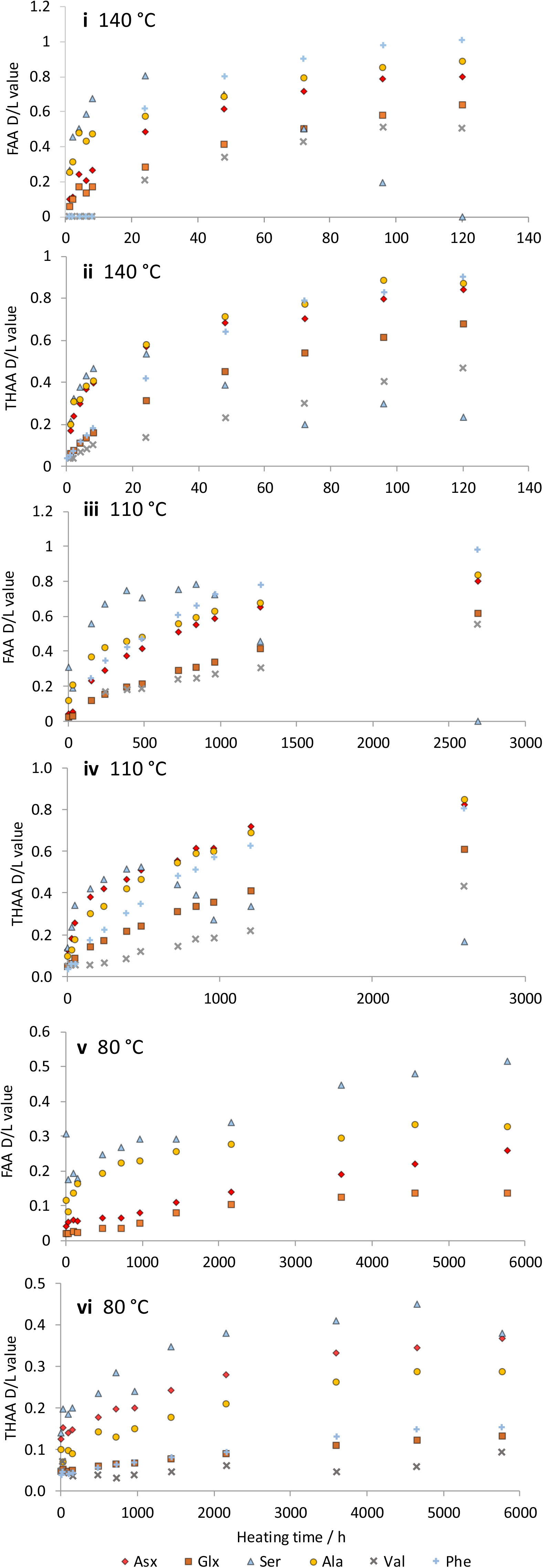
*Extent of racemization of FAAs and THAAs in the intra-crystalline fraction of enamel during 140 (i & ii), 110 (iii & iv) and 80 (v & vi) °C elevated temperature experiments*.

#### 5.5.1 Calculation of reaction parameters using elevated-temperature data

The use of AAR as a tool for absolute age estimation requires the relationship between the age and extent of racemization to be described. This can be achieved through mathematically modelling accelerated degradation experiments, coupled with calibration using direct dating (e.g. radiometric dating) and/or indirect geochronological evidence from the same geographic region (Miller *et al*, 1983; Murray-Wallace & Bournman, 1990; Wehmiller, 2013). Methods using high temperature experimental data assume that the mechanisms for racemization *in situ* are synonymous with the process occurring at elevated temperatures, which is not the case for all biominerals (Tomiak *et al.*, 2013). To mathematically describe the kinetic behaviour of enamel proteins, two models have been tested: reversible first order rate kinetics (RFOK) and constrained power law kinetics (CPK; Clarke and Murray-Wallace, 2006).

#### 5.5.2 Reversible first order reaction kinetics (RFOK) to describe racemization in enamel

Racemization of FAAs is theoretically described by reversible first-order kinetics (RFOK), assuming no thermal decomposition of amino acids or additional side reactions (Bada & Schroeder, 1972; Shou & Bada, 1980). This behaviour has been shown to predict the racemization of aqueous FAAs relatively well (Bada and Schroeder, 1975; Smith and Reddy, 1989), but does not always accurately describe the intra-crystalline racemization of amino acids in biominerals (Penkman *et al.*, 2008; Crisp *et al.*, 2013; Tomiak *et al.*, 2013). This is not unexpected, as the complexity of the inter- and intra-molecular interactions involved in the degradation of peptide-bound amino acids in biominerals would be highly unlikely to follow RFOK (Collins & Riley, 2000; Clarke & Murray-Wallace, 2006).

RFOK dictates that:

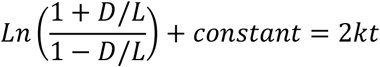

*Equation 1. k is the rate of racemization and is equal in both the forward and backward reaction, t is time and n is varied to improve the description of the relationship between D/L and time*.

If ln{(1+D/L)/(1-D/L)} has a linear relationship with time then the mechanism for racemization can be described by RFOK. The strength of the correlation has been assessed based on coefficient of determination values (R^2^). To model RFOK in fossil biominerals, Crisp *et al.* (2013) suggested excluding data that yielded R^2^ values < 0.97, as it was not appropriate to linearize the data below this grade of correlation. When including the full range of time points for enamel, the R^2^ values for most of the plots was < 0.97, even when restricted ranges of D/L were used (Table 5). The amino acids with the highest conformity to RFOK were Glx and Phe (Table 5; Figure 17). Val also yielded correlations higher than the 0.97 threshold for heating experiments run at 110 and 140 °C, but not for 80 °C. The rate of racemization of Val in enamel is very slow and thus the extent of racemization observed with in the time frame of the 80 °C experiment was not long enough to acquire a high degree of correlation.

The plot of ln[(1+D/L)/(1-D/L)] vs. t for Asx exhibits two D/L ranges which can be linearised with dissimilar gradients and therefore kinetic behaviours (Figure 17 and Table 5). This lack of conformity to RFOK is likely to be a consequence of the complex nature of Asx racemization, which is a composite signal encompassing both aspartic acid and asparagine (Glx is also a composite signal).

**Table 5.**
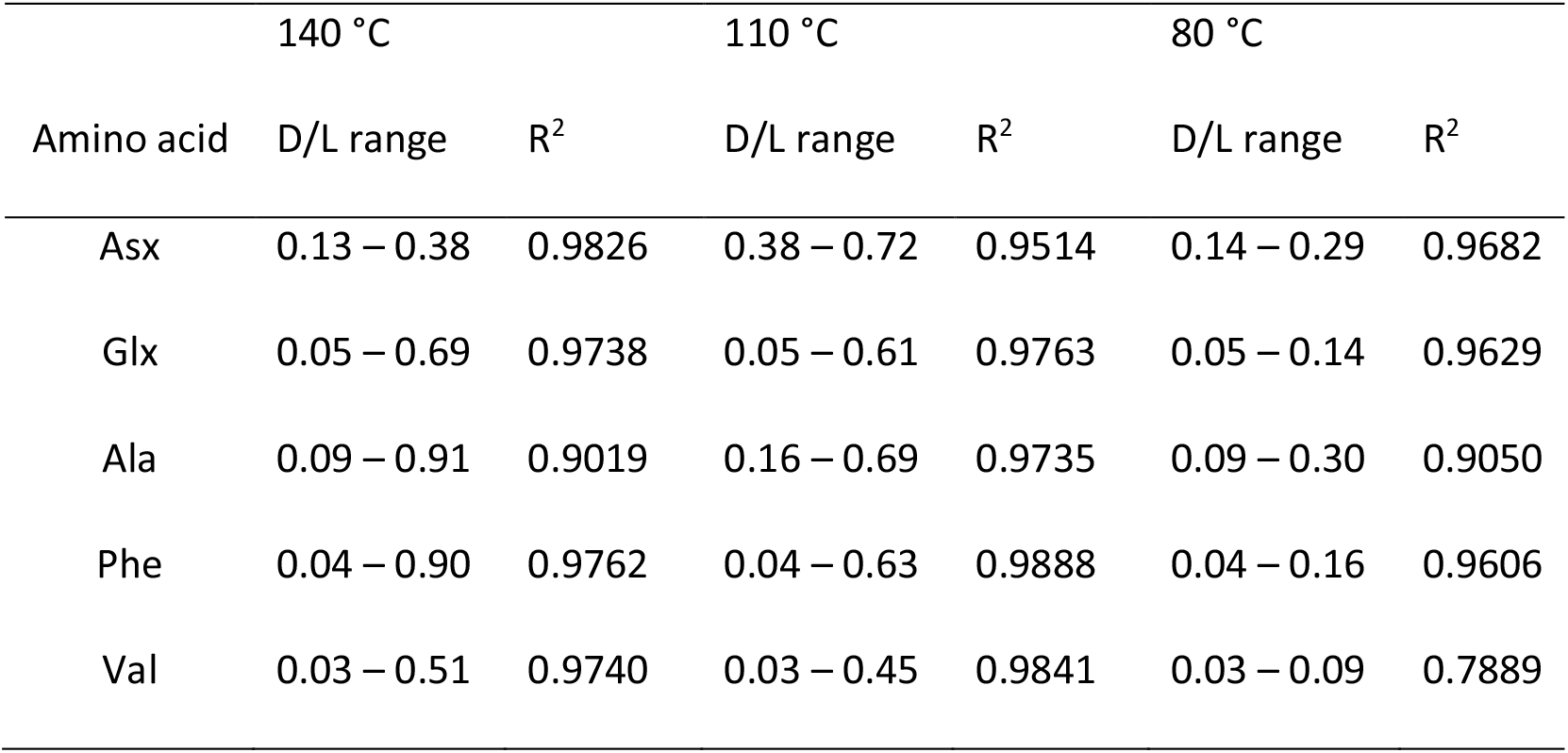
*R^2^ values from the plot between the transformed D/L values against time, constrained D/L ranges have been used to optimise correlation. R^2^ values test for conformity to RFOK*.

| Amino acid | 140 °C |  | 110 °C |  | 80 °C |  |
| --- | --- | --- | --- | --- | --- | --- |
| | D/L range | $R^2$ | D/L range | $R^2$ | D/L range | $R^2$ |
| Asx | 0.13 – 0.38 | 0.9826 | 0.38 – 0.72 | 0.9514 | 0.14 – 0.29 | 0.9682 |
| Glx | 0.05 – 0.69 | 0.9738 | 0.05 – 0.61 | 0.9763 | 0.05 – 0.14 | 0.9629 |
| Ala | 0.09 – 0.91 | 0.9019 | 0.16 – 0.69 | 0.9735 | 0.09 – 0.30 | 0.9050 |
| Phe | 0.04 – 0.90 | 0.9762 | 0.04 – 0.63 | 0.9888 | 0.04 – 0.16 | 0.9606 |
| Val | 0.03 – 0.51 | 0.9740 | 0.03 – 0.45 | 0.9841 | 0.03 – 0.09 | 0.7889 |

**Figure 17.**
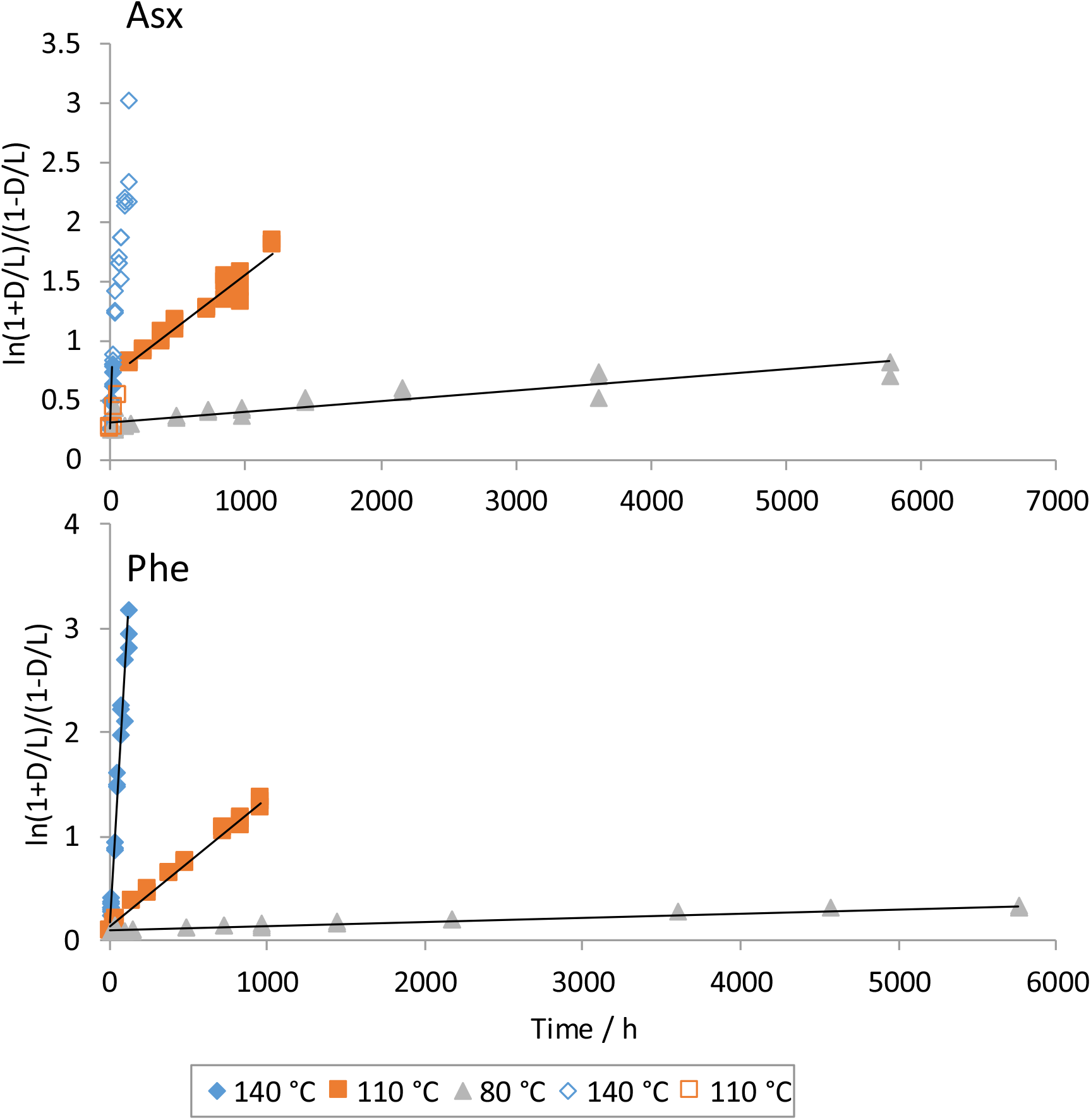
*Transformed D/L data on increasing heating time at three different temperatures. Linearity of the plot indicates adherence to RFOK; an R^2^ cut-off of 0.97 is used in this study. The gradient is proportional to the rate of reaction. Block colour data points were used to calculate rates and assess correlation. Outlined data points were not used to calculate the rates of reactions as they reduced the strength of the correlation*.

#### 5.5.3 Constrained power-law kinetics (CPK) to describe racemization in enamel

The lack of adherence between RFOK and heating experiments in other biominerals has led to the use of alternative, experimentally derived functions to describe amino acid racemization (e.g. Manley *et al.*, 2000; Kaufman *et al.*, 2000; Clarke & Wallace 2006; Allen *et al.*, 2013; Crisp *et al.*, 2013; Tomiak *et al.*, 2013). Constrained power-law kinetics (CPK) can be used to improve the description of a trend by raising one of the terms to a power (Equation 2). This power (n) is varied to maximise the R^2^ value and thus improve the mathematical description of the trend (e.g. Figure 18). CPK has been used with the most success to better describe the racemization of Asx and Glx (Kaufman, 2000, Manley *et al.*, 2000; Crisp *et al.*, 2013).

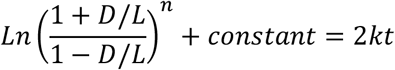

*Equation* 2: k *is the rate of racemization and is equal in both the forward and backward reaction*, t *is time and n is varied to improve the description of the relationship between D/L and time*.

The use of CPK can better describe the rate of racemization in enamel for most of the amino acids at a specific temperature (Table 6). However, for most amino acids there is little concordance between the optimised power values the transformed D/L value are raised to at each of the different temperatures. For example, the optimum value of n to maximise the linear fit of Phe using CPK at 140 and 110 °C is 1.2 and for 80 °C is 1.7 (Figure 18). This indicates that across different D/L value ranges/temperatures the data are exhibiting a different relationship between transformed D/L value and time. This lack of coherence between temperatures indicates that trend fitting in this way may not be appropriate for most amino acids in enamel.

**Table 6.** *R^2^ values from the plot between the transformed D/L values against time; constrained D/L ranges have been used to optimise correlation. R^2^ values test for conformity to CPK at the specified value of n. Values in bold signify the highest R^2^ for that temperature and specific amino acid*.

| Amino acid | n | 140 °C |  | 110 °C |  | 80 °C |  |
| --- | --- | --- | --- | --- | --- | --- | --- |
| | | D/L range | $R^2$ | D/L range | $R^2$ | D/L range | $R^2$ |
| Asx | 0.3 | 0.13 – 0.38 | 0.9569 | 0.38 – 0.72 | **0.9555** | 0.14 – 0.29 | 0.9496 |
| Asx | 1.5 | 0.13 – 0.38 | **0.9881** | 0.38 – 0.72 | 0.9326 | 0.14 – 0.29 | 0.9745 |
| Asx | 1.8 | 0.13 – 0.38 | 0.9863 | 0.38 – 0.72 | 0.9205 | 0.14 – 0.29 | **0.9755** |
| Glx | 1.7 | 0.05 – 0.69 | **0.9972** | 0.05 – 0.61 | **0.9972** | 0.05 – 0.17 | **0.9787** |
| Ala | 1.1 | 0.09 – 0.95 | **0.9140** | 0.16 – 0.69 | 0.9779 | 0.09 – 0.30 | 0.9117 |
| Ala | 1.5 | 0.09 – 0.95 | 0.9434 | 0.16 – 0.69 | **0.9858** | 0.09 – 0.30 | 0.9329 |
| Ala | 2.2 | 0.09 – 0.95 | 0.9441 | 0.16 – 0.69 | 0.9699 | 0.09 – 0.30 | **0.9479** |
| Phe | 1.3 | 0.04 – 0.90 | **0.9816** | 0.04 – 0.63 | **0.9945** | 0.04 – 0.16 | 0.9728 |
| Phe | 1.7 | 0.04 – 0.90 | 0.9689 | 0.04 – 0.63 | 0.9849 | 0.04 – 0.16 | **0.9780** |

**Figure 18.**
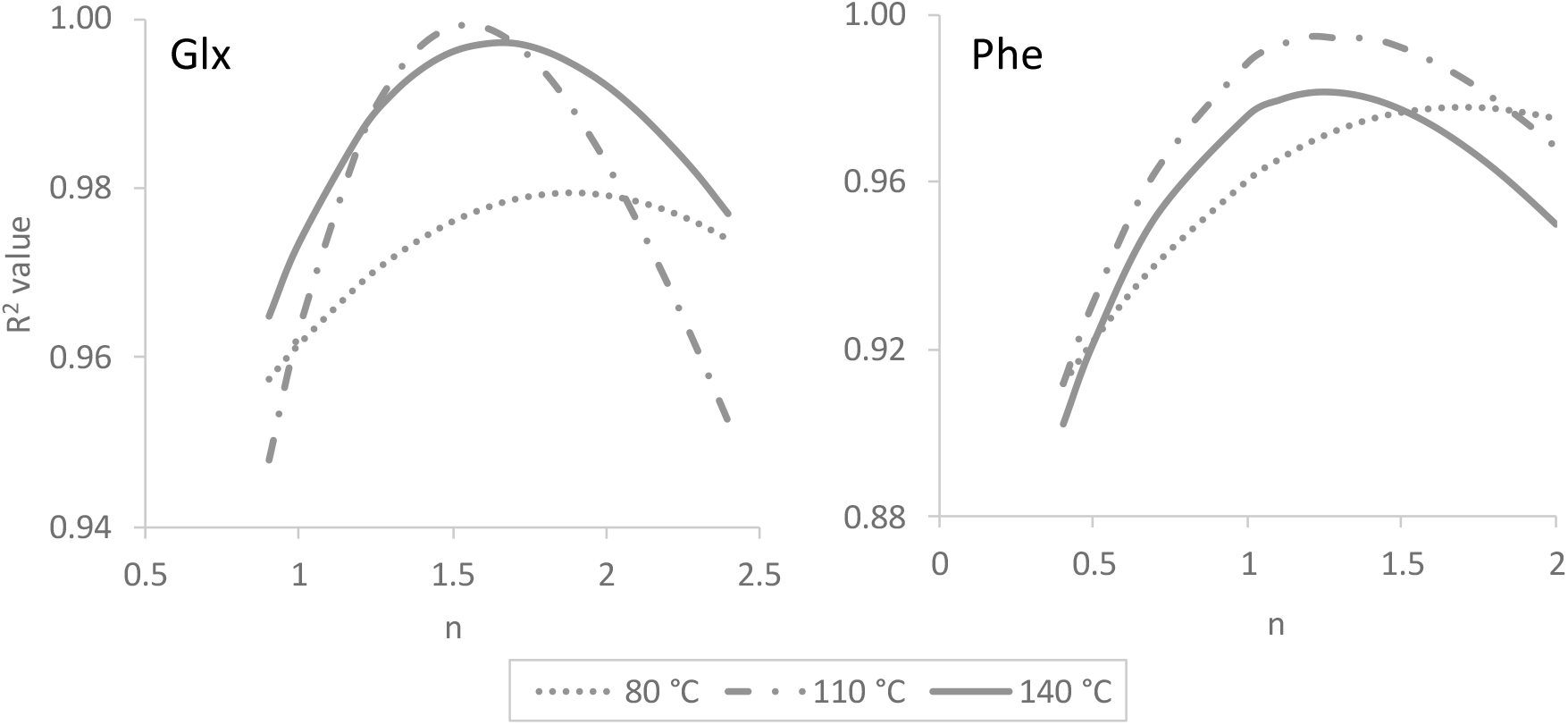
*R^2^ values for the three different temperature heating experiments when D/L values are transformed using CPK. R^2^ values have been plotted against the power (n) ln{(1+D/L)/(1-D/L)} has been raised to (Equation 2)*.

#### 5.5.4 Arrhenius parameters

The calculated rate constants at the various temperatures can be used to determine kinetic parameters using the Arrhenius equation (Equation 3). This calculation is reliant on accurate calculated rates of racemization and assumes a consistent mechanism for racemization across the studied range of D/L values and temperatures.

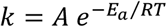

*Equation* 3*: k is the rate constant, A is the frequency factor, E_a_ is the activation energy, R is the gas constant and T is the temperature*.

**Table 7.** *Activation energies, frequency factors (A) and Arrhenius plot R^2^ values for the four studied amino acids. Ranges and R^2^ values for the RFOK and CPK data used are shown in Table 5 & Table 5 respectively*.

| Amino acid | Model | Arrhenius plot $R^2$ value | $E_a$ / kJ mol<sup>-1</sup> | ln($A$) |
| --- | --- | --- | --- | --- |
| Asx | RFOK | 0.9790 | 121 | 25.7 |
| Glx | RFOK | 0.9984 | 123 | 30.7 |
| Glx | CPK | 0.9999 | 143 | 37.1 |
| Ala | RFOK | 0.9882 | 109 | 27.1 |
| Phe | RFOK | 0.9998 | 128 | 33.1 |

In contrast to the other amino acids studied, the R^2^ values for Glx can roughly be optimised using CPK (n = 1.7), yielding a good linear relationship between the transformed D/L values and heating time between 0.05 – 0.69 D/L values (Table 6). Some of the kinetic parameters of the degradation of Glx have therefore been calculated and methods for trend fitting have been compared to assess the potential for using high temperature studies to estimate absolute age and palaeotemperature reconstruction with enamel AAR.

The activation energy for racemisation for Glx calculated using a CPK method for ostrich eggshell was 147 kJ mol^−1^ (Crisp *et al.*, 2013) and 141 kJ mol^−1^ *for Patella vulgata* (Demarchi *et al.*, 2013c), compared to 123 kJ mol^−1^ using RFOK and 143 kJ mol^−1^ using CPK for enamel. The activation energies for different biominerals are likely to be diverse, so it is not surprising that the value calculated for enamel would be different to those of ostrich eggshell and *Patella vulgata*. The divergence in the values between the models for enamel are significant however; it is therefore difficult to assess the true value for the activation energy.

#### 5.5.5 Palaeothermometry using Glx

The relationship between rate of racemization and ambient temperature of the reaction medium (T_eff_) can be extrapolated to the burial environment provided the kinetics of the racemization reaction and temperature sensitivity of the rate constants are known (Miller *et al.*, 1983; Oches *et al.*, 1996). To highlight the sensitivity and importance of the choice of mathematical model for estimating kinetic parameters, 13 proboscidean enamel samples with well constrained temporal assignments from the UK, plus a modern Asian elephant tooth (*E. maximus*), have been used to estimate the ambient temperature of the reaction medium (T_eff_) in the UK over the last 2 Ma using RFOK (Figure 19) and CPK. Using RFOK, T_eff_ is calculated to be −10 °C and using a CPK model T_eff_ is 1 °C (Figure 20). These predicted temperatures are unlikely; comparable experimental data and palaeoclimatic data indicate ranges of 8-10 °C are more realistic (BGS report, 2011). Not only is there a significant discrepancy between the temperatures obtained using the different models, the absolute values indicate that the assumption of similar kinetics over these temperature ranges may not be valid.

**Figure 19.**
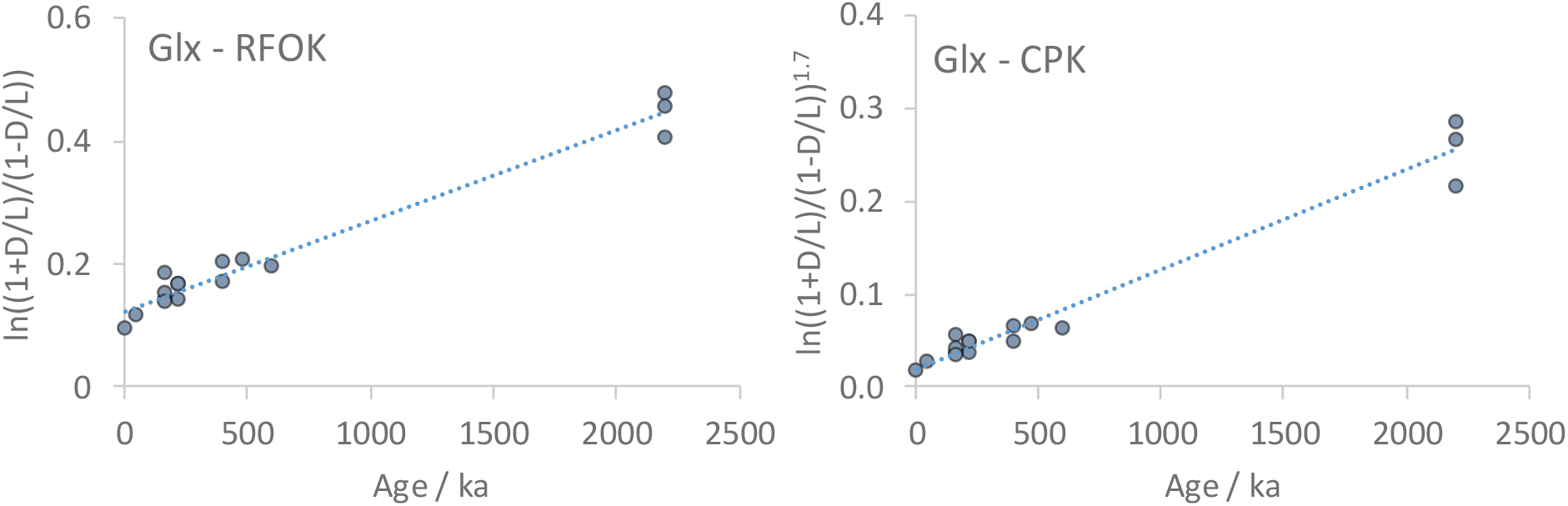
*Linearised relationship between age of fossil enamel from UK sites and transformed D/L value assuming RFOK (left) and CPK (right). Gradient of the liner trend is proportional to the rate of racemization*.

**Figure 20.**
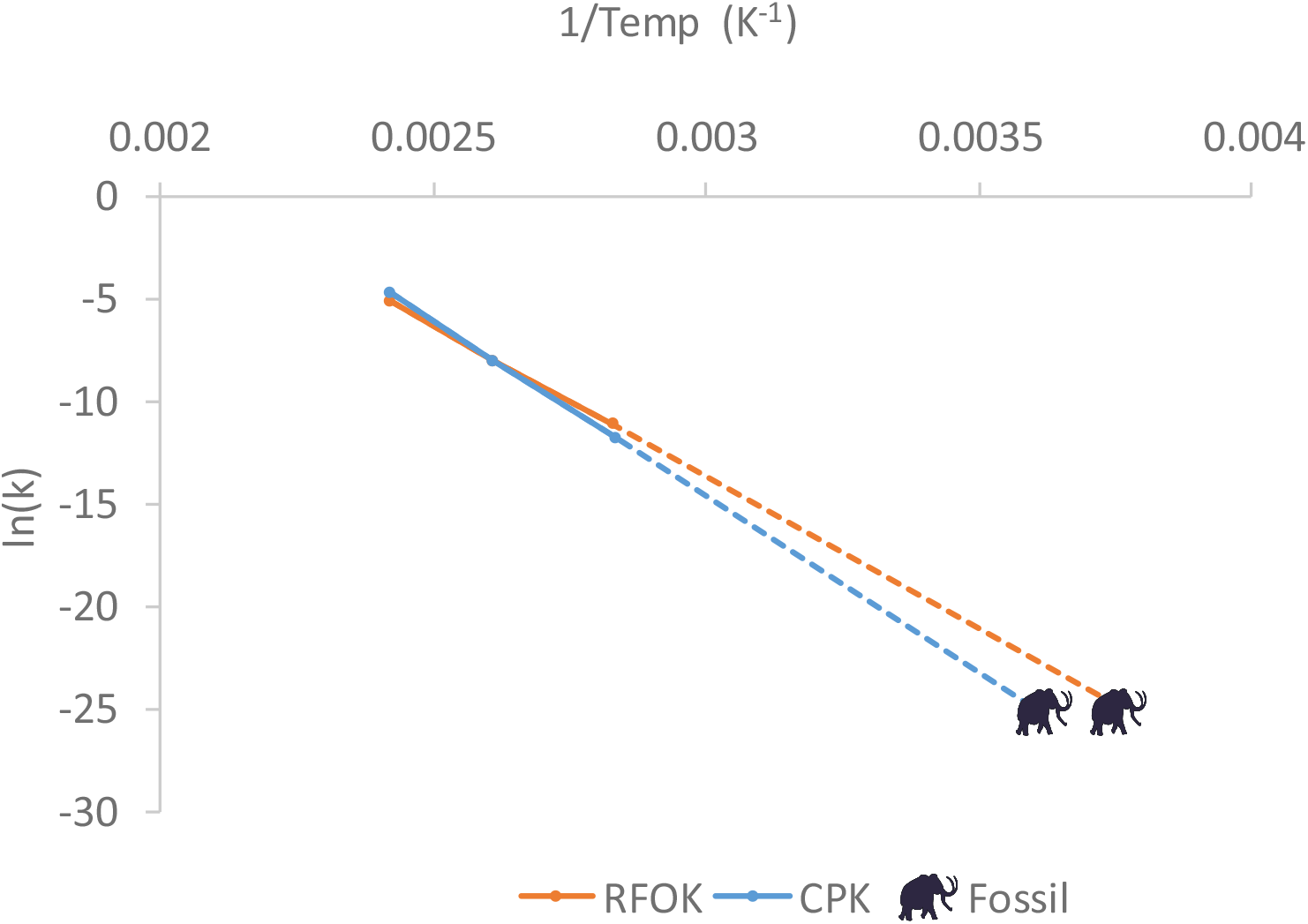
*Arrhenius plot used to calculate the activation energies using RFOK and CPK. Effective temperatures of the fossil UK samples have been calculated using each of the models using the rate of reaction of enamel fossil material*.

#### 5.5.6 Enamel fossil data and comparison to *Bithynia* opercula heated experiments

To understand the kinetics in more detail, the patterns of racemization in enamel were compared to those obtained from *Bithynia* opercula, using both fossil samples and elevated temperature experimental data. The extent of racemization in fossil proboscidean enamel increases with independent evidence of age but is considerably lower than in *Bithynia* opercula when compared to samples from the same deposits (Figure 21). The lower rates of racemisation in enamel (cf. *Bithynia*) suggest that the enamel AAR may be able to be used as a relative dating technique over longer time scales than *Bithynia*. However, this will likely be balanced by a lower temporal resolution due to the slower rates yielding smaller incremental changes over time.

The rates of racemization are very similar between the two biominerals in the elevated-temperature experiments (Figure 22). This indicates that the rates of racemization in enamel and opercula are not behaving in comparable ways at elevated temperatures and in the depositional environment. The lack of consistency between the two data sets indicates these elevated temperature experiments do not fully describe the degradation mechanisms in the depositional environment for enamel. If the rates of reaction calculated in elevated temperatures are not appropriate for extrapolation to the burial environment, then irrespective of how accurately a mathematical model is describing those rates, derivation of absolute age estimates using the kinetic parameters calculated in this manuscript is therefore inappropriate.

**Figure 21.**
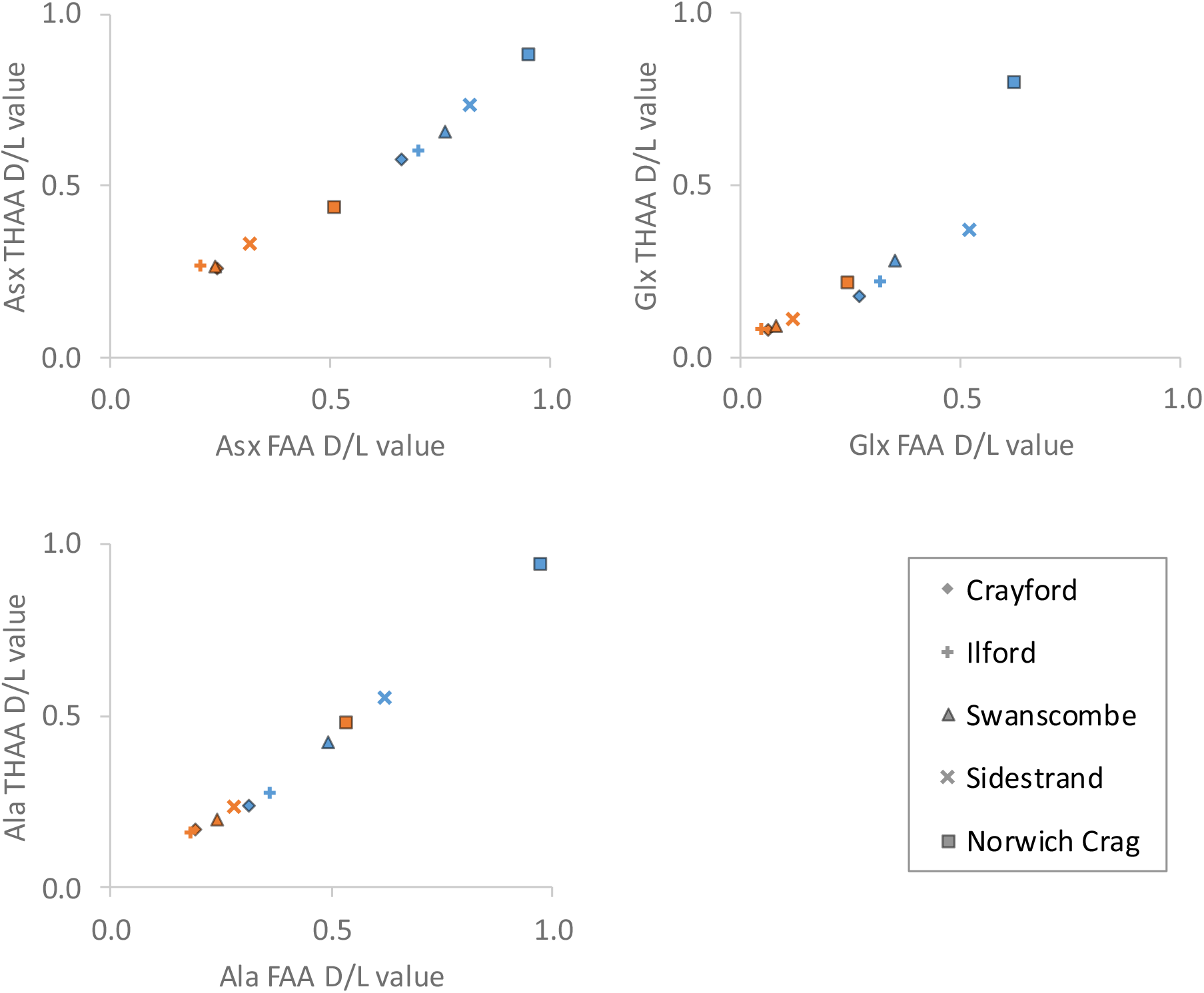
*Extent of racemization in fossil enamel (orange) and* Bithynia *opercula (blue) IcPD from a range of UK sites. Sites have been selected to give a broad range of ages spanning the Quaternary*.

**Figure 22.**
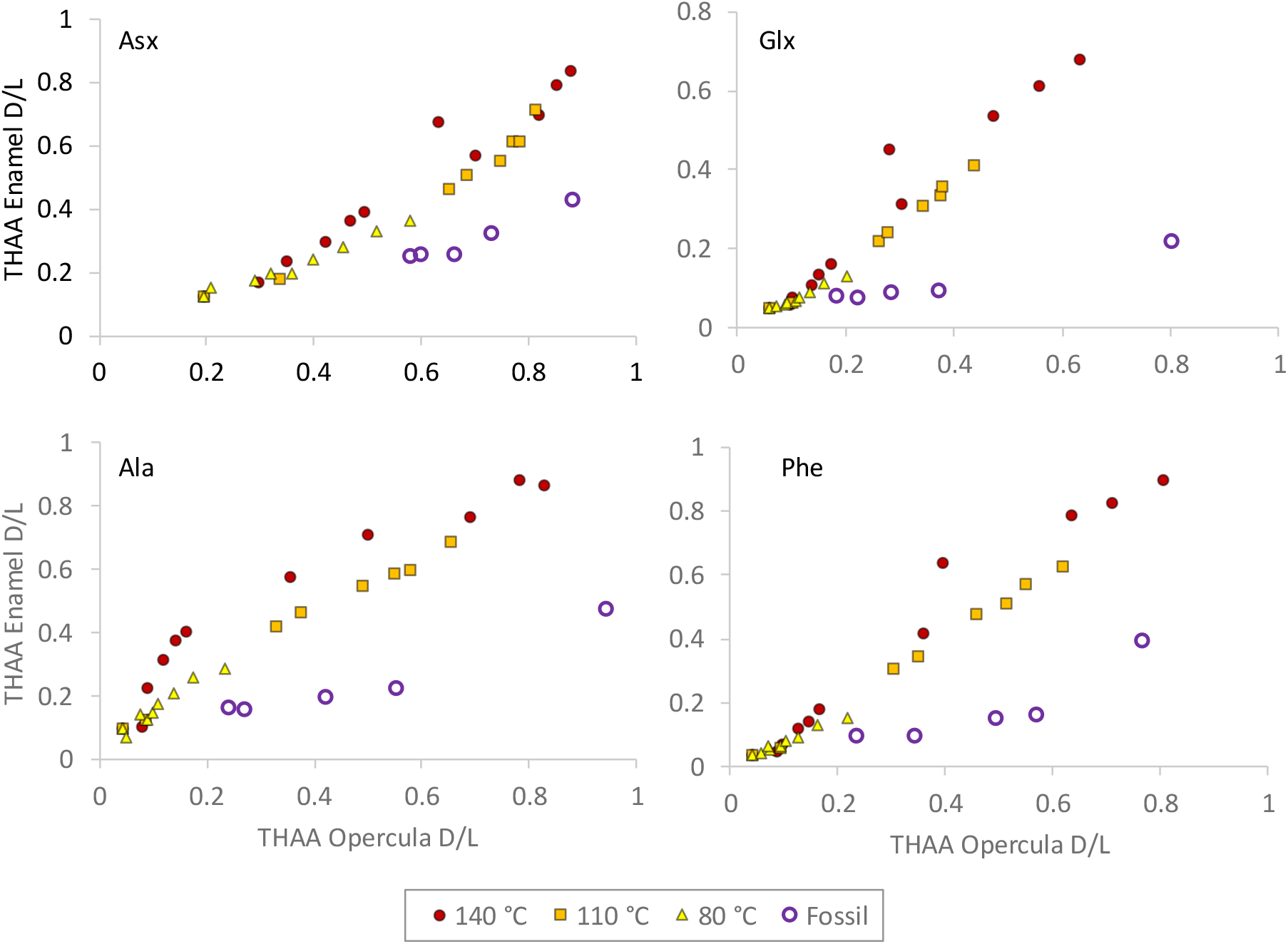
*Extents of THAA racemization in proboscidean enamel plotted against* Bithynia *opercula values for both elevated temperature experiments at 140, 110 and 80 °C (blue, orange and grey respectively), and for fossil data from sites where both opercula and enamel could be compared. Opercula elevated temperature data points have been plotted against their equivalent time point from the enamel study (at the same temperature)*.

## 6 Conclusions

A simple protocol for amino acid analysis of enamel has been developed, involving a biphasic separation of the inorganic species (originating from the mineral matrix) from the amino acids. This enables the routine analysis of fossil enamel proteins via a RP-HPLC method. Crucially the method does not greatly influence the amino acid composition and does not alter the amino acid D/L values.

A fraction of amino acids, stable to prolonged exposure to a strong oxidant has been isolated (ca. 72 h exposure to NaOCl) and has been shown to exhibit effectively closed system behaviour during simulated degradation experiments. Only minimal leaching of amino acids into the supernatant water was observed during diagenetic experiments (ca. 5%), and at least part of that total is due to the presence of small crystals of enamel suspended in solution.

Elevated temperature studies have shown that enamel intra-crystalline amino acids follow predictable trends of racemization and breakdown, but the divergence with patterns observed in fossil enamel indicate that different mechanisms are prevailing at lower diagenetic temperatures. This therefore limits the ability to use high temperature experiments to calculate relevant kinetic parameters. However, the predictability of the diagenetic behaviour of enamel amino acids in both the fossil and elevated temperature experiments, indicate that analysis of the intra-crystalline protein fraction of enamel is suitable for relative amino acid geochronology over Quaternary timescales.

In conclusion, we present here a new preparative method for the routine analysis of closed system amino acids from fossil enamel. This technique has the potential to build geochronologies for time scales on the order of millions of years and therefore to directly estimate the age of material where there previously has not been an applicable technique.

## 7 Acknowledgements

We acknowledge support from NERC (NE/K500987/1 and NE/L501761/1) and the Leverhulme Trust (PLP-2012-116) for providing funding for the project. We also acknowledge the help and support from Paul Shepherd and the staff at the British Geological Survey for access to their collections for sampling, and for technical support from Sheila Taylor and Oliver Bayfield.

## References

Akiyama M., 1980. Diagenetic decomposition of peptide-linked serine residues in the fossil scallop shells. In: Biogeochemistry of amino acids. Hare, P. E., Hoering T. C., King K., 115–120. John Wiley & Sons.

Allen A. P., Kosnik M. A., Kaufman D. S., 2013. Characterizing the dynamics of amino acid racemization using time-dependent reaction kinetics: A Bayesian approach to fitting age-calibration models. Quaternary Geochronology 18, 63–77.

Amend J. P., Helgeson, H. C, 1997. Solubilities of the common L-a-amino acids as a function of temperature and solution pH. Pure and applied chemistry 69, 935–942.

Ashton N. 2018. Landscapes of habit and persistent places during MIS 11 in Europe. In: Pope, M., McNabb J., Gamble C, (eds). Crossing the Human Threshold: Dynamic Transformation and Persistent Places during the Middle Pleistocene, London: Routledge, 142–164.

Bada J. L. 1985. Amino acids racemization dating of fossil bones. Annual Review Earth Planetary Science 13, 241–268.

Bada J. L & Schroeder R. A., 1975. Amino acid racemization reactions and their geochemical implications. Die Naturwissenschaften 62, 71–79.

BGS Report, 2011. Geo Reports. BGS Repot No: GR_000000/1.

Bischoff J. L & Rosenbauer R. J., 1981. Uranium series dating of human skeletal remains from the del Mar and Sunnyvale sites, California. Science 213, 1003–1005.

Blackwell B., Rutter N. W., Debénath A. 1990. Amino acid racemization in mammalian bones and teeth from La Chaise‐de‐Vouthon (Charente), France. Geoarchaeology 5, 121–147.

Brandon A. & Sumbler M. G., 1991. The Balderton Sand and Gravel: Pre-Ipswichian cold stage fluvial deposits near Lincoln, England. Journal of Quaternary Science 6, 117–138.

Bravenec A. D., Ward K. D., Ward T. J., 2018. Amino acid racemization and its relation to geochronology and archaeometry. Separation Science 41, 1489–1506.

Bridgland D. R., 1994. Quaternary of the Thames. (Eds.) Chapman & Hall, London.

Bridgland D.R., Gibbard P. L., Harding P., Kemp R. A., Southgate G., 1985. New information and results from recent excavations at Barnfield pit, Swanscombe. Quaternary Newsletter 46, 25–39.

Bridgland D.R., Howard A.J., White M.J., White T.S., 2014. The Quaternary of the Trent. Oxbow Books, Oxford.

Bridgland D. R., Howard A. J., White M. J., White T. S., Westaway R., 2015. New insight into the Quaternary evolution of the River Trent, UK. Proceedings of the Geologists’ Association 126, 466–479.

Bright J. & Kaufman D. S., 2011. Amino acid racemization in lacustrine ostracodes, part I: effect of oxidizing pre-treatments on amino acid composition. Quaternary Geochronology 6, 154–173.

Brooks A. S., Hare P. E., Kokis J. E., Miller G. H., Ernst R. D., Wendorf F., 1990. Dating Pleistocene Archeological Sites by Protein Diagenesis in Ostrich Eggshell. American Association for the Advancement of Science 248, 60–64.

Canoira L., García-Martínez, M.J., Llamas J. F., Ortíz J. E., Torres T., 2003. Kinetics of Amino Acid Racemization (Epimerization) in the Dentine of Fossil and Modern Bear Teeth. International Journal of Chemical Kinetics 35, 576–591.

Chandler R. H., 1914. The Pleistocene Deposits of Crayford. Proceeding of the Geological Association 25, 61–71.

Clarke S. J., Murray-Wallace, C.V., 2006. Mathematical expressions used in amino acid racemization geochronology: a review. Quaternary Geochronology 1, 261–278.

Collins M. J. & Riley M. S., 2000. Amino acid racemization in biominerals, the impact of protein degradation and loss. In: Goodfriend, G. A., Collins M. J., Fogel M. L., Macko S. A., Wehmiller J. F. (Eds.), Perspectives in Amino Acid and Protein Geochemistry. Oxford University Press, 120–142.

Collins M. J., Waite E. R., van Duin, A C., 1999. Predicting protein decomposition: the case of aspartic-acid racemization kinetics. Philosophical Transactions of the Royal Society B Biological Sciences 354, 51–64.

Collins M. J., Penkman K. E. H., Rohland N., Shapiro B., Dobberstein R. C., Ritz-Timme S., Hofreiter M., 2009. Is amino acid racemization a useful tool for screening for ancient DNA in bone? Proceedings of the Royal society B 276. 1–7.

Coope G. R., 2001. Biostratigraphical distinction of interglacial coleopteran assemblages from southern Britain attributed to Oxygen Isotope Stages 5e and 7. Quaternary Science Reviews 20, 1717–1722.

Crenshaw M. A., 1972. The soluble matrix from *Mercenaria mercenaria* shell. Biomineralisation 6, 6–11.

Crisp M., Demarchi B., Collins M., Morgan-Williams M., Pilgrim E., Penkman K., 2013. Isolation of the intra-crystalline proteins and kinetic studies in *Struthio camelus* (ostrich) eggshell for amino acid geochronology. Quaternary Geochronology 16, 110–128.

Demarchi B., Rogers, K. Fa, D. A., Finlayson C. J., Milner N., Penkman K. E. H., 2013a. Intra-crystalline protein diagenesis (IcPD) in *Patella vulgata*. Part I: Isolation and testing of the closed system. Quaternary Geochronology 16, 144–157.

Demarchi B., Collins M., Bergström E., Dowle A., Penkman K., Thomas-Oates J., Wilson J., 2013b. New experimental evidence for in-chain amino acid racemization of serine in a model peptide. Analytical Chemistry 85, 5835–5842.

Demarchi B., Collins M. J., Tomiak P. J., Davies B. J., Penkman K. E. H., 2013c. Intra-crystalline protein diagenesis (IcPD) in *Patella vulgata*. Part II: breakdown and temperature sensitivity. Quaternary Geochronology 16, 158–172.

Dirks P. H. G. M., Roberts, Eric M., Hilbert-Wolf H., Kramers J. D., Hawks J., Dosseto A., Duval M., Elliott M., Evans M., Grun R., Hellstrom J., Herries A. I. R., Joannes-Boyau R., Makhubela T. V., Placzek C. J., Robbins J., Spandler C., Wiersma J., Woodhead J., Berger L. R., 2017. The age of *homo naledi* and associated sediments in the rising star cave, South Africa. eLife e24231.

Duval M., 2015. Evaluating the accuracy of ESR dose determination of pseudo-Early Pleistocene fossil tooth enamel samples using dose recovery tests. Radiation Measurements 79, 24–32.

Frouin M., Lahaye C., Valladas H., Higham T., Debénath A., Delagnes A., 2017. Dating the Middle Paleolithic deposits of La Quina Amont (Charente, France) using luminescence methods. Journal of Human Evolution 109, 30–45.

Garot E., Couture-Veschambre C., Manton D., Rodriguez V., Lefrais Y., Rouas P., 2007. Diagnostic guide enabling distinction between taphonomic stains and enamel hypomineralisation in an archaeological context. Archives of Oral Biology 74, 28–36.

Gash A. E., Tillotson T. M., Satcher J. H., Hrubesh, L W., Simpson R. L., 2001. New sol-gel synthetic route to transition and main-group metal oxide aerogels using inorganic salt precursors. Journal of Non-Crystalline Solids 285, 22–28.

Girling M. A., 1974. Evidence from Lincolnshire of the age and intensity of the Mid-Devensian temperate episode. Nature 250, 270.

Gries K., Kröger R., Kübel C., Fritz M., Rosenauer A., 2009. Investigations of voids in the aragonite platelets of nacre. Acta Biomaterialia 5, 3038–3044.

Griffin R., 2006. Application of amino acid racemization in enamel to the age estimation of age at death of archaeological remains. PhD Thesis.

Grün R., 1991. Appendix 5. Electron spin resonance age estimates on elephant teeth from the Balderton Sand and Gravel. In: Brandon, A. & Sumbler M. G. The Balderton Sand and Gravel: Pre-Ipswichian cold stage fluvial deposits near Lincoln, England. Journal of Quaternary Science 6, 135–136.

Grün R., Aubert M., Hellstrom J., Duval M., 2010. The challenge of direct dating old human fossils. Quaternary International 223-224, 87-93.

Hamblin R. J. O., Moorlock B. S. P., Booth S. J., Jeffery D. H., Morigi A. N., 1997. The Red Crag and Norwich Crag formations in eastern Suffolk. Proceedings of the Geologists’ Association 108, 11–23.

Hare P. E., 1988. Organic geochemistry of bone and its relation to the survival of bone in the natural environment. In: Behrensmeyer, A. K. & Hill A. P., Fossils in the making: Vertebrate taphonomy and paleoecology. The University of Chicago Press.

Hare P. E. & Mitterer R.M., 1967. Non-protein amino acids in fossil shells. Carnegie Institution of Washington 65, 362–364.

Hendy E. J., Tomiak P. J., Collins M. J., Hellstrom J., Tudhope A. W., Lough J. M., Penkman K. E. H., 2012. Assessing amino acid racemization variability in coral intra-crystalline protein for geochronological applications. Geochimica et Cosmochimica Acta 86, 338–353.

Hershkovitz I., Weber G. W., Quam R., Duval M., Grun R., Kinsley L., Ayalon A., Bar-Mattews M., Valladas H., Mercier N., Arsuaga J. L., Martinon-Torres M., Bermudez de Castro, J. M., Fornai C., Martin-Frances L., Sarig R., May H., Krenn V. A., Slon V., Rodriguez L., Garcia R., Lorenzo C., Carretero J. M., Frumkin A., Shahack-Gross R., Mayer D. E. B., Cui Y., Wu X., Peled N., Groman-Yaroslavski I., Weissbrod L., Yeshurun R., Tsatskin A., Zaidner Y., Weinstein-Evron M., 2018. The earliest modern humans outside Africa. Science 359, 456–459.

Holyoak D. T. and Preece R. C., 1985. Late Pleistocene Interglacial deposits at Tattershall, Lincolnshire. Philosophical Transactions of the Royal Society of London B, Biological Sciences 311, 1149, 193–236.

Ingalls A. E., Lee C., Druffel E. R. M., 2003. Preservation of organic matter in mound-forming coral skeletons. Geochimica et Cosmochimica Acta 67, 2827–2841.

Jacobi R. M., Higham T. F G., Bronk R. C., 2006. AMS radiocarbon dating of Middle and Upper Palaeolithic bone in the British Isles: Improved reliability using ultrafiltration. Journal of Quaternary Science 21, 557–573.

Kaufman D. S., 2000. Amino acid racemization in ostracodes. In: Goodfriend, G. A., Collins M. J., Fogel M. L., Macko S. A., Wehmiller J. F. (Eds.), Perspectives in Amino Acid and Protein Geochemistry. Oxford University Press, New York, 145–160.

Kaufman D. S. and Manley W. F., 1998. A new procedure for determining DL amino acid ratios in fossils using reverse phase liquid chromatography. Quaternary Science Reviews 17, 987–1000.

King W.B.R. and Oakley K. P., 1936. The Pleistocene Succession in the Lower parts of the Thames Valley. Proceedings of the Prehistoric Society 2, 52–76.

Kriausakul N. and Mitterer R. M., 1978. Isoleucine epimerization in peptides and proteins: kinetic factors and application to fossil proteins. Science 201, 1011–1014.

Lister A. M., 1993. The stratigraphical significance of deer species in the Cromer forest‐bed formation. Journal of Quaternary Science 8, 95–108.

Lister A. M., 1996. The stratigraphical interpretation of large mammal remains from the Cromer Forest-bed Formation. In: The Early Middle Pleistocene in Europe ed. 25–44. Rotterdam.

Lister A. M., 1998. The age of Early Plistocene mammal faunas from the ‘Weybourne Crag’ and Cromer Forest-bed Formation (Norfolk, England). Mededelingen Nederlands Instituut voor Toegepaste Geowetenschappen 60, 271–280.

Lister A. M. and Brandon A., 1991. A pre-Ipswichian cold stage mammalian fauna from the Balderton Sand and Gravel, Lincolnshire, England. Journal of Quaternary Science 6, 139–157.

MacPhee R. D. E., Tikhonov A. N., Mol D., de Marliave C., van der Plicht, H., Greenwood A. D., 2002. Radiocarbon Chronologies and Extinction Dynamics of the Late Quaternary Mammalian Megafauna of the Taimyr Peninsula, Russian Federation. Journal of Archaeological Science 29, 1017–1042.

Manley W. F., Miller G. H., Czywczynski J., 2000. Kinetics of aspartic acid racemization in Mya and Hiatella: modelling age and palaeotemperature of high-latitude Quaternary molluscs. In: Goodfriend, G. A., Collins M. J., Fogel M. L., Macko S. A., Wehmiller J. F. (Eds.), Perspectives in Amino Acid and Protein Geochemistry. Oxford University Press, New York, 120–141.

Margolis H. C., Beniash E., Fowler C. E., 2006. Role of macromolecular assembly of enamel matrix proteins in enamel formation. Journal of Dentistry Research 85, 775–793.

Marin F., Luquet G., Marie B., Medakovic D., 2008. Molluscan Shell Proteins: Primary Structure, Origin, and Evolution. Current Topics in Developmental Biology 80, 209–276.

Marshall E., 1990. Racemization dating: great expectations. Science 247, 799.

Meijer T. & Preece R. C., 2000. A review of the occurrence of *Corbicula* in the Pleistocene of NorthWest Europe. Geologie en Mijnbouw/Netherlands Journal of Geosciences 79, 241–255.

Miller G. H., Sejrup H. P., Mangerud J., Andersen B. G., 1983. Amino acid ratios in Quaternary molluscs and foraminifera from western Norway: correlation, geochronology and paleotemperature estimates. Boreas 12, 107–124.

Mitterer R. M., Kriausakul N., 1984. Comparison of the rates and degrees of isoleucine epimerization in dipeptides and tripeptides. Organic Geochemistry 7, 91–98.

Mol D., Post K., Reumer J. W. F., van der Plicht, J., de Vos J., van Geel, van Reenen, G., Pals J. P., Glimmerveen J., 2006. The Eurogeul - First report of the palaeontological, palynological and archaeological investigations of this part of the North Sea. Quaternary International 142-143, 178-185.

Murray-Wallace C. V., Bourman R. P., 1990. Direct radiocarbon calibration for amino acid racemization dating. Australian Journal of Earth Sciences 37, 365–367.

Oches E. A., McCoy W. D., Clark P. U., 1996. Amino acid estimates of latitudinal temperature gradients and geochronology of loess deposition during the last glacial maximum, Mississippi Valley, United States. Geological Society of America Bulletin 108, 892–903.

Orem C. A., Kaufman D. S., 2011. Effects of basic pH on amino acid racemization and leaching in freshwater mollusk shell. Quaternary Geochronology 6, 233–245.

Ortiz J. E., Sánchez-Palencia Y., Gutiérrez-Zugasti, I., Torres T., González-Morales M., 2018. Protein diagenesis in archaeological gastropod shells and the suitability of this material for amino acid racemization dating: *Phorcus lineatus* (da Costa, 1778). Quaternary Geochronology 46, 16–27.

Paz A., Guadarrama D., López M., González J. E., Brizuela N., Aragón J., 2012. A comparative study of hydroxyapatite nanoparticles synthesized by different Routes. Química Nova 35, 1724–1727.

Penkman K. E. H., Preece R. C., Keen D. H., Maddy D., Schreve D. C., Collins M. J., 2007. Testing the aminostratigraphy of fluvial archives: the evidence from intra-crystalline proteins within freshwater shells. Quaternary Science Reviews 26, 2958–2969.

Penkman K. E. H., Kaufman D. S., Maddy D., Collins M. J., 2008. Closed-system behaviour of the intra-crystalline fraction of amino acids in mollusc shells. Quaternary Geochronology 3, 2–25.

Penkman K. E. H., Preece R. C., Bridgland D. R., Keen D. H., Meijer T., Parfitt S. A., White T. S., Collins M. J., 2011. A chronological framework for the British Quaternary based on *Bithynia* opercula. Nature 476, 446–449.

Penkman K. E. H., Preece R. C., Bridgland D. R., Keen D. H., Meijer T., Parfitt S. A., White T. S., Collins M. J., 2013. An aminostratigraphy for the British Quaternary based on *Bithynia* opercula. Quaternary Science Reviews 61, 111–134.

Poinar H. N., Stankiewicz B. A., 1999. Protein preservation and DNA retrieval from ancient tissues. Proceedings of the National Academy of Sciences of the United States of America 96, 8426–8431.

Powell J., Collins M. J., Cussens J., Macleod N., Penkman K. E. H., 2013. Quaternary. Geochronology Results from an amino acid racemization inter-laboratory proficiency study; design and performance evaluation. Quaternary Geochronology 16, 183–197.

Preece R. C., 1999. Mollusca from Last Interglacial fluvial deposits of the River Thames at Trafalgar Square, London. Journal of Quaternary Science 14, 77–89.

Preece R. C. & Penkman K. E. H., 2005. New faunal analyses and amino acid dating of the Lower Palaeolithic site at East Farm, Barnham, Suffolk. Proceedings of the Geologists’ Association 116, 363–377.

Preece R. C., Parfitt S. A., Coope G. R., Penkman K. E. H., Ponel P., Whittaker J. E., 2009. Biostratigraphic and aminostratigraphic constraints on the age of the Middle Pleistocene glacial succession in north Norfolk, UK. Journal of Quaternary Science 24, 557–580.

Rackman D. J., 1978. Evidence for the changing vertebrate communities in the Middle Devensian. Quaternary Newsletter 25, 1–3.

Refsnider K. A., Miller G. H., Frechette B., Rood D. H., 2013. A chronological framework for the Clyde Foreland Formation, Eastern Canadian Arctic, derived from amino acid racemization and cosmogonic radionuclides. Quaternary Geochronology 16, 21–34.

Robinson C., Kirkham J., Brookes S. J., Bonass W. A., Shore R. C., 1995. The chemistry of enamel development. International Journal of Developmental Biology 39, 145–152.

Robinson C. Brookes, S. J., Shore R. C., Kirkham J., 1998. The developing enamel matrix: nature and function. Oral sciences 106, 282–291.

Schreve D. C., 2001. Differentiation of the British late Middle Pleistocene interglacials: the evidence from mammalian biostratigraphy. Quaternary Science Reviews 20, 1693–1705.

Shou P. M., & Bada J. L. 1980. The pK’s of Amino Acids at Elevated Temperatures Estimated from Racemization Data. Naturwissenschaften 67, 37–38.

Smith G. G. and Evans R. C., 1980. The effect of structure and conditions on the rate of racemisation of free and bound amino acids. In P.E. Hare, T.C. Hoering, and K. King Jr., Eds., Biogeochemistry of Amino Acids, 257–282, New York: Wiley.

Smith G. G. & Reddy G. V., 1989. Effect of the side-chain on the racemization of amino-acids in aqueous-solution. Journal of Organic Chemistry 54, 4529–4535.

Stephenson R. C. and Clarke S., 1989. Succinimide formation from aspartyl and asparaginyl peptides as a model for the spontaneous degradation of proteins. Journal of Biological Chemistry 264, 6164–6170.

Stuart A. J. 1982. Pleistocene Vertebrates in the British Isles. London: Longman.

Sutcliffe A. J., 1975. A hazard in the interpretation of glacial - interglacial sequences. Quaternary Newsletter 17, 1–3.

Sykes G. A., Collins M. J., Walton D. I., 1995. The significance of a geochemically isolated intracrystalline organic fraction within biominerals. Organic Geochemistry 23, 1059–1065.

Takahashi O., Kobayashi K., Oda A., 2010. Computational insight into the mechanism of serine residue racemization. Chemistry and Biodiversity 7, 1625–1629.

Taylor R. E., 1983. Non-concordance of Radiocarbon and Amino Acid Racemization Deduced Age Estimates on Human Bone. Radiocarbon 25, 647–654.

Tomiak P. J., Penkman K. E H., Hendy E. J., Demarchi B., Murrells S., Davis S., McCullagh P., Collins M. J., 2013. Testing the limitations of artificial protein degradation kinetics using known-age massive Porites coral skeletons. Quaternary Geochronology 16, 87–109.

Towe K. M., 1980. Preserved organic ultrastructure: An unreliable indicator for Paleozoic amino acid biogeochemistry. Hare, P. E., Hoering T. C., King K. (Eds.), Biogeochemistry of Amino Acids. Wiley, New York, pp. 65–74.

Towe K. M. and Thompson G. R., 1972. The structure of some bivalve shell carbonates prepared by ion-beam thinning - A Comparison Study. Calcified Tissue Research 10, 38–48.

Turner-Walker G., 2007. The chemical and microbial degradation of bones and teeth. Advances in human palaeopathology (eds). John Wiley & Sons, Ltd. 11–15.

Vasenko, L &, Qu H., 2018. Calcium phosphates recovery from digester supernatant by fast precipitation and recrystallization. Journal of Crystal Growth 481, 1–6.

Wehmiller J. F., 2013. Interlaboratory comparison of amino acid enantiomeric ratios in Pleistocene fossils. Quaternary Geochronology 16, 173–182.

Wehmiller J. F., Harris W. B., Boutin B. S., Farrell K. M., 2012. Calibration of amino acid racemization (AAR) kinetics in United States mid-Atlantic Coastal Plain Quaternary mollusks using 87Sr/86Sr analyses: Evaluation of kinetic models and estimation of regional Late Pleistocene temperature history. Quaternary Geochronology 7, 21–36.

Zalasiewicz J. A., Mathers S. J., Gibbard P. L., Peglar S. M., Funnell B. M., Catt J. A., Harland R., Long P. E., Austin T. J. F., 1991. Age and Relationships of the Chillesford Clay (Early Pleistocene: Suffolk, England). Philosophical Transactions of the Royal Society B: Biological Sciences 333, 81–100.

